# Optogenetic control of actin crosslinker length reveals a mechanical basis for cortical symmetry breaking

**DOI:** 10.64898/2026.08.30.748082

**Authors:** Filipe Nunes Vicente, Anumita Jawahar, Michael Wassermair, Mahsa Rahimi, Aliaksandr Dzementsei, Martin Kräter, Lena Fischer, Rubén Tesoro-Moreno, Baptiste Vauleon, Jochen Guck, Anđela Šarić, Ivan Palaia, Matthieu Piel, Olivia du Roure, Julien Heuvingh, Alba Diz-Muñoz

## Abstract

Cell shape changes during migration, division, or differentiation require the dynamic regulation of actin network mechanics. Actin crosslinkers are central to this regulation, controlling network connectivity and the transmission of contractile forces. A large diversity of crosslinkers exists, differing in length, domain structure, and binding kinetics, yet why cells deploy specific crosslinkers in a physiological context remains unclear.

To bridge this gap, we developed a light-controlled actin crosslinker toolbox spanning three physiologically relevant lengths: ∼9 nm (fascin-like), ∼16 nm (fimbrin-like), and ∼56 nm (alpha-actinin-like). Using magnetic pincher experiments and *in silico* modelling, we show that short and mid-length crosslinkers dynamically tune cortical stiffness and thickness in a density- and myosin-dependent manner, with short crosslinkers also driving pronounced stress-stiffening as the cortex is deformed. Strikingly, minute-scale activation reveals a length-dependent switch in cell behaviour: short crosslinkers cause cortical delamination, while long ones instead drive cell polarization and symmetry breaking. This switch can be overridden by perturbing actin turnover, which unlocks polarization in mid-length crosslinkers that otherwise delaminate. Crosslinker-induced polarization is not merely a local cortical event: it directs subsequent cell spreading, coupling a nanometre-scale molecular choice to a cell-scale decision about movement.

Together, these findings establish a versatile optogenetic platform for manipulating actin crosslinking, and show that the cortex can encode a behavioural switch directly in its material architecture.

## INTRODUCTION

Cell shape changes underlie many of the most fundamental processes in biology, from single-cell events like polarization, division, and migration to multicellular processes such as morphogenesis and wound healing^1–3^. In animal cells, these changes are largely driven by the cell cortex, a thin, dynamic network of actin filaments and myosin motors that lies beneath the plasma membrane^2,3^. The cortex is built and regulated by more than 100 actin-binding proteins (ABPs), which control the nucleation, branching, severing, membrane attachment, and crosslinking of actin filaments^4^. To generate the forces driving these shape changes, myosin binds to actin filaments and slides them past one another at the expense of ATP hydrolysis, resulting in cortical tension^1,5^. In addition to myosin activity, actin crosslinkers are crucial to ensure force propagation, generating networks with different architectures and material properties. These, in turn, determine how the network responds to myosin-generated stress and thus its contractile behavior^4,6^. The cortex contains a large diversity of actin crosslinkers, including fascin, plastin, alpha- actinin, filamin, and spectrin^3^, which can regulate a broad spectrum of biological processes, from cortical polarization and cytokinetic ring formation during cell division^7–9^, to force sensing at epithelial junctions^10^. These crosslinkers have evolved over long timescales across the eukaryotic tree of life and exhibit diverse structural properties as well as actin-binding domains^11,12^. For example, crosslinkers of the spectrin superfamily, including alpha-actinin, contain two calponin-homology (CH) actin-binding domains separated by tandem arrays known as spectrin repeats^13^; fascin, on the other hand, belongs to an entirely different branch and binds actin through a unique β-trefoil configuration^14^. Given this considerable diversity, how and when cells choose which crosslinker to employ for a specific process remains an unanswered question.

Over the last decade, several *in vitro* studies have focused on how crosslinker density regulates the material properties of the network and its force transmission. In reconstituted actin networks, long- range coordinated contraction only occurs above a percolation threshold of crosslinker density, at which all actin filaments become connected and stress can propagate across the entire system^15,16^. Below this threshold, filaments are loosely interconnected and only contract at short scales, forming asters^15,16^. Under compression, individual actin filaments can also buckle, causing networks to soften rather than stiffen^17^. At high crosslinker concentrations, by contrast, networks become progressively stiffer upon increasing deformation, a nonlinear behavior known as strain-stiffening^18^.

Beyond concentration, the physical and biochemical properties of individual crosslinkers also shape network architecture and mechanics. Molecular length sets inter-filament spacing: short crosslinkers like fascin and plastin (∼5–6 nm^14,19^ and ∼9.3–9.8 nm^20^, respectively) form tightly bundled networks with filaments spaced only ∼8–15 nm apart^14,19,20^, although the structural flexibility of fascin enables it to form hexagonal arrays^14^. Longer crosslinkers like alpha-actinin (∼30–36 nm^21^) produce more widely spaced bundles (∼35–40 nm^22,23^), while the much longer filamin (∼40–160 nm^24,25^) and spectrin (∼90–180 nm^26,27^), generate meshworks spanning entire subcellular compartments^26–28^. Length also affects network dynamics: filamin-generated meshworks exhibit greater mechanical hysteresis than those formed by the shorter alpha-actinin^29^, potentially reflecting greater flexibility and rearrangement in longer crosslinkers. Moreover, crosslinkers can segregate into networks of different inter-filament distances due to their intrinsic crosslinker properties^23,30^. Beyond length, crosslinker binding-site geometry can also determine filament polarity: fascin and fimbrin, for instance, generate unipolar and mixed-polarity bundles, respectively^31^, which in turn shapes myosin motor dynamics and force generation within the bundle^32,33^.

This diversity in crosslinker properties is matched by the diversity in their deployment: cells target specific crosslinkers to specific structures, matching molecular properties to functional needs. Short crosslinkers such as fascin and filamin populate fast-remodeling structures like the leading edge of migrating cells and filopodia^30,32^, while longer ones such as alpha-actinin and fimbrin dominate in more stable structures, such as stress fibers and microvilli^34,35^. At the cortex, plastin limits cortical polarization in the *C. elegans* embryo, while a developmental switch between fimbrin and filamin reorganizes hexagonal actin arrays during *Drosophila* cellularization^36^. Similar specialization segregates distinct spectrin isoforms to axons versus dendrites in neurons^37,38^. This picture is further complicated by cooperativity: multiple crosslinkers often act together within the same network, further shaping its architecture and mechanical behaviour in ways that cannot be predicted from any single crosslinker’s properties alone^7^. Together, these examples show that their activity must be precisely coordinated in space and time to support cellular structures and sustain global force transmission. Yet why cells deploy specific ones for specific processes remains poorly understood, as no system has so far allowed crosslinking to be manipulated with the precision cells themselves achieve over space, time, length, and density.

In this study, we engineer a scalable, optogenetic toolbox of actin crosslinkers to control cortical mechanics and dynamics over second-to-minute timescales. Using magnetic pincher measurements, we show that crosslinker activation increases cortical stiffness and thickness in a length- and myosin-dependent manner, revealing how the molecular properties of actin crosslinkers can dynamically tune the macroscopic properties of the cell surface. We then show that long crosslinkers trigger cortical polarization and symmetry breaking, while short crosslinkers instead promote cortical delamination — both processes regulated by myosin-driven contractility. We further show that perturbing actin turnover can unlock polarization in intermediate-length crosslinkers that otherwise cause delamination. Finally, we show that this crosslinker-induced symmetry breaking directs cell spreading. Together, these results position crosslinker length as a tunable dial that cells can use to switch between distinct cortical behaviors, linking a purely physical network parameter to cell-scale decisions such as polarization and motility.

## RESULTS

### Developing a scalable optogenetic actin crosslinker toolbox

Molecular optogenetics is a powerful approach to control cell motility, polarization, and contractility over space and time by fusing light-sensitive molecular switches to effector proteins such as RhoA^39^, ROCK modulators^40^, and nonmuscle myosin II^41^. To achieve spatial and temporal control of actin crosslinking, we engineered optogenetic actin crosslinkers containing the blue light-responsive cryptochrome 2 (CRY2), fused to the Utrophin CH1&2 (UtCH) actin-binding domain and an mCherry tag (UtCH-CRY2) (**Fig. 1A**). Crosslinking of these proteins is activated via CRY2 homo-oligomerization upon illumination with blue light (488 nm)^42,43^ (**Fig. 1A**).

**Figure 1.**
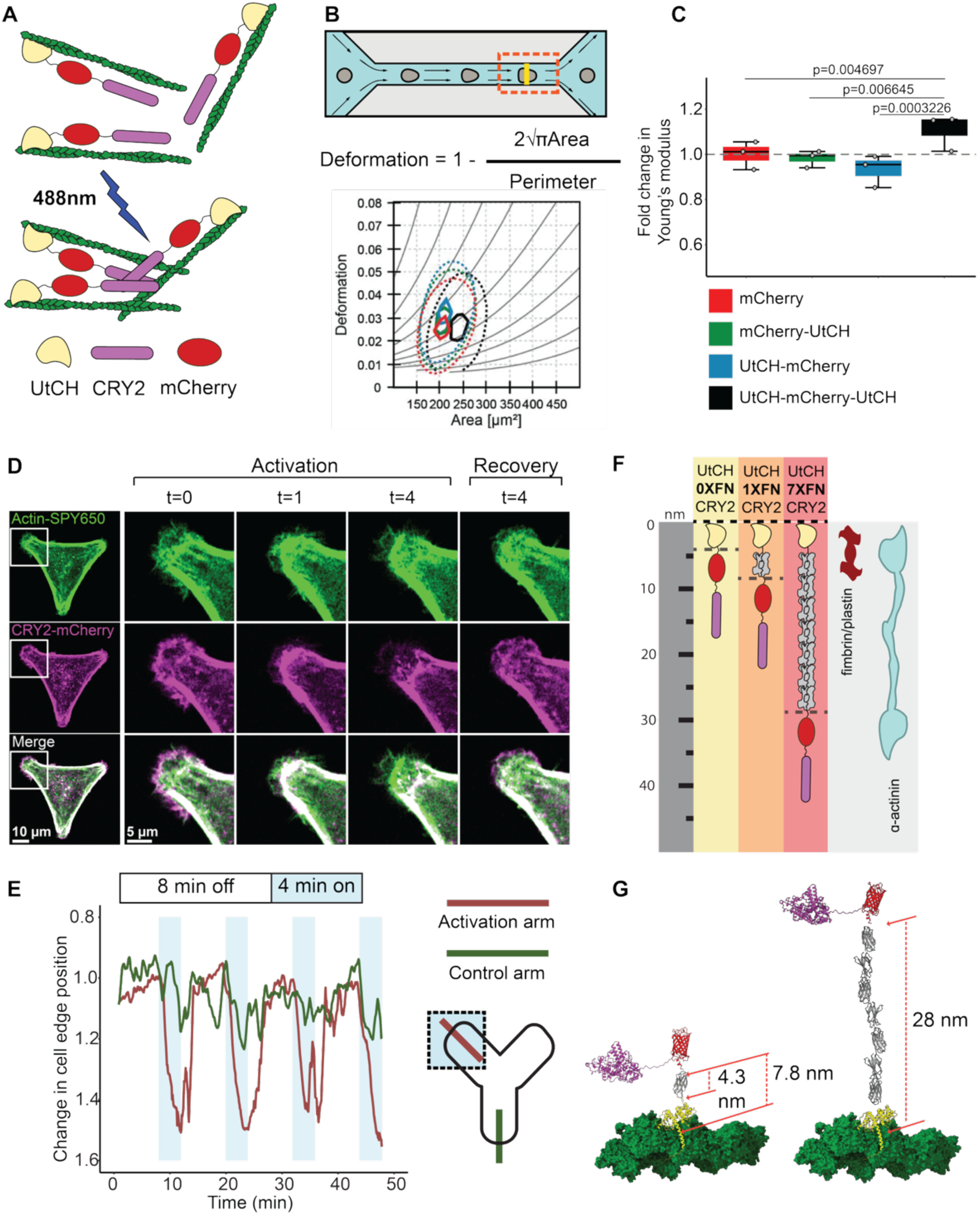
An optogenetic toolbox to control actin crosslinking. **A)** Schematic representation of an optogenetic actin crosslinker. A utrophin-CH domain is combined with CRY2 and tagged with mCherry. Upon blue light (488nm) illumination, the optogenetic crosslinker drives oligomerization of actin filaments. **B)** (**top**) RTDC setup. (**bottom**) Microfluidics deformation and area measurements of 3T3 fibroblasts expressing either UtCH-mCherry-UtrCH (black), mCherry-UtrCH (green), UtrCH-mCherry (blue) or mCherry (red) as a control. Isolasticity lines (grey) indicate trends in stiffness. Contour plots were generated using multivariate kernel density estimation. Solid lines correspond 50% and dashed lines correspond to 95% of events. **C)** Young’s modulus for RTDC experiments was derived from deformation and area measurements using numerical simulation for fully elastic spheres according to Mokbel et al., 2017^50^. Boxplots represent average mean values, normalized to mCherry average, from 3 independent experiments. Error bars represent standard deviation between experiments. Statistical analysis using 3 independent measurements was performed with Linear mixed models. **D)** Airyscan images of 3T3 fibroblasts expressing 0X-CRY2 (magenta) with live actin FastAct- SPY650 staining (green) on a Y-shaped micropattern during an activation cycle followed by a dark phase. **(left)** Airyscan image of the full cell with the activation region in white. **(middle and right)** Zoom of the highlighted region during activation and recovery cycles. **E)** Position of cell edge over time for multiple activation (4 min, blue shade) and recovery (8 min, white) cycles for activated region (dark red) and a control region of the cell (green). Higher values correspond to cell edge retraction. **F)** To increase the molecular length of the crosslinker, a Fibcon domain (grey) is used. Schematic of 0X(yellow), 1XFibcon (1XFN, orange) and 7XFibcon (7XFN, red) optogenetic crosslinkers with predicted lengths of 3.5 nm (0X), 7.8 nm (1X), and 28nm (7X) and endogenous crosslinkers of similar lengths to 1XFN and 7XFN. **G)** Representative AlphaFold3 models edited and superimposed with ChimeraX to display UtCH-1XFibcon-CRY2 (left) and UtCH-7XFibcon-CRY2 (right) crosslinkers bound to 4 beta-actin chains. Beta-actin (green), UtCH (yellow), Fibcon (grey), mCherry (red), CRY2 (magenta). For UtCH-1X-Fibcon-CRY2, 7.8 nm distance was calculated between the centre of mass of binding site at beta-actin and the center of mass of 1XFibcon; 4.3 nm distance was calculated between the centre of mass of UtCH and 1XFibcon. For UtCH-7X-Fibcon-CRY2, 28 nm distance was calculated between calculated between the centre of mass of binding site at beta-actin and the centre of mass of the last Fibcon in the 7X spacer. See also Figure S1.

**Figure S1.**
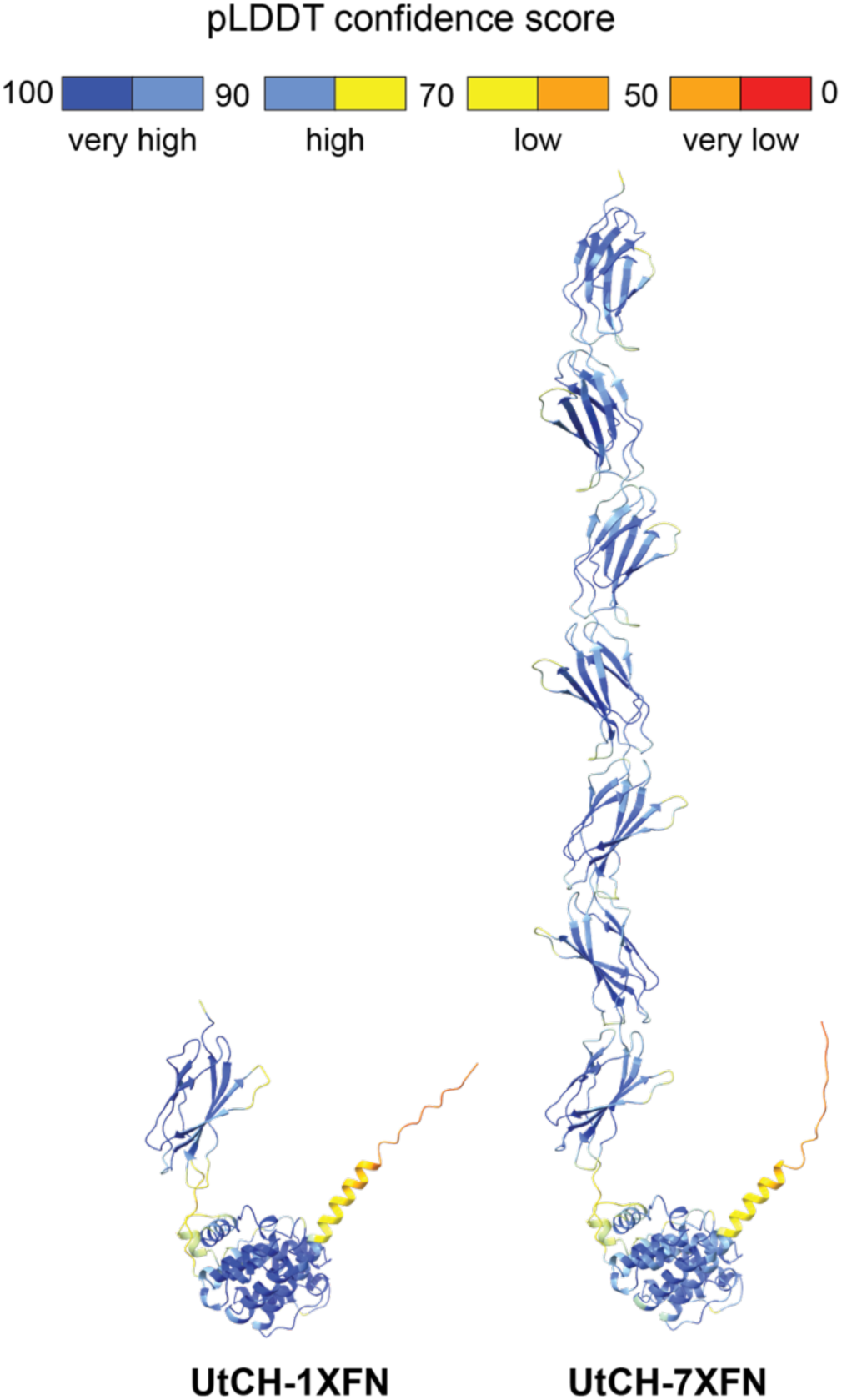
1XFN and 7XFN form a rigid spacer for actin crosslinkers. Selected views of the predicted AlphaFold3 model for 1XFN or 7XFN bound to UtCH, coloured for predicted local distance difference test (pLDDT) values.

To understand whether this approach could induce a mechanical phenotype, we first generated a constitutive bivalent actin crosslinker, UtCH-mCherry-UtCH, in which two UtCH domains flank an mCherry tag, allowing it to crosslink actin filaments independently of light activation. UtCH-mCherry and mCherry- UtCH, each containing a single UtCH domain, served as monovalent, non-crosslinking controls. All constructs were expressed in a doxycycline-inducible manner in NIH-3T3 fibroblasts. We then performed real-time deformability cytometry (RT-DC), which allows high-throughput measurement of mechanical properties^44^. In RT-DC, cells are pumped through a microfluidic channel and imaged by bright-field microscopy. In the narrow constriction zone, cells deform due to pressure gradients and shear stress, and deformation is determined by image processing^44^ (**Fig. 1B**). Expression of UtCH-mCherry-UtCH produced an increase of cell elasticity compared to UtCH-mCherry and mCherry-UtCH controls, confirming that bivalent actin crosslinking via UtCH is sufficient to increase cell stiffness (**Fig. 1C**).

Next, we tested our light-inducible construct, UtCH-CRY2, to determine whether its activation could modulate actin dynamics in a spatiotemporal manner. For that, we combined live high-resolution imaging of 3T3 mouse fibroblasts in Y-shaped micropatterns with cycles of subcellular activation with blue light (**Fig. 1D, E**). Subcellular activation of UtCH-CRY2 at actin protrusions triggered protrusion retraction in a reversible and repetitive manner, while the control region exhibited consistent dynamics across multiple activation cycles (**Fig. 1E**). Therefore, UtCH-CRY2 can regulate actin dynamics in live cells over space and time in a reversible manner.

We next generated a toolbox encompassing optogenetic crosslinkers of different molecular lengths and expressed them in a doxycycline-inducible manner in NIH-3T3 fibroblasts. To achieve this, we harnessed a class of protein domains known as Fibcons^45^, designed as the structural consensus of 15 natural fibronectin FN3 domains with high stability. Individual Fibcons can be sequentially combined to form long, semi-flexible polymers^46^. Fibcon-based polymers have been used to develop surface crowding sensors^47^ and modulate antigen height^48^, making them ideal candidates to increase crosslinker length in a scalable manner. Therefore, we inserted two distinct Fibcon assemblies (1XFibcon or 7XFibcon) into our original optogenetic crosslinker and generated two novel constructs with different lengths: UtCH- 1XFibcon-CRY2 (1X-CRY2) and UtCH-7XFibcon-CRY2 (7X-CRY2) (**Fig. 1F**). The original crosslinker is henceforth referred to as 0X-CRY2.

To estimate the length of the optogenetic crosslinkers, we used AlphaFold3^49^ to predict the structure of 1XFN and 7XFN bound to UtCH. The resulting models exhibited high confidence metrics (**Fig. S1**). Moreover, the models reproducibly displayed the length of 1 Fibcon domain as ∼4.3nm and the structure of 7XFN as a long, rigid chain of stacked Fibcons (**Fig. S1**), in line with previously quantified distances *in vitro*^46^. We defined crosslinker length as the distance from the UtCH actin-binding site to the last Fibcon residue before mCherry (or, for 0X-CRY2, which lacks a Fibcon domain, directly from the predicted position of the UtCH actin-binding site). This yielded lengths of ∼3.5 nm for 0X-CRY2 (similar to fascin at ∼5-6 nm^14,19^), ∼7.8 nm for 1X-CRY2 (similar to fimbrin/plastin at ∼9.3-9.8 nm^20^), and ∼28 nm for 7X-CRY2 (close to alpha-actinin at ∼30-36 nm^21^) (**Fig. 1G**). Because our synthetic crosslinkers dimerize through CRY2, unlike endogenous crosslinkers, the resulting inter-filament distance can be up to two-fold larger than the crosslinker’s own length. Therefore, the inter-filament distance would be 7 nm for 0X-CRY2 (similar to fascin’s, measured at ∼8.1-12 nm^14,19^), 15.6 nm for 1X-CRY2 (still similar to fimbrin/plastin’s, measured at 14.6-14.9 nm^20^) and 56 nm for 7X-CRY2 (larger than alpha-actinin’s, measured at 35-40 nm^22^).

### Actin crosslinkers regulate cortical mechanics in a length-dependent manner

The mechanical properties of the cortex, in particular stiffness and tension, are the primary determinant of cellular mechanics and morphology^2^. Cortical architecture and network connectivity have emerged as key parameters in force transmission and regulation of contractility, shaped by both crosslinking^16,32,51^ and actin filament length^52^. However, how the dynamic rearrangement of actin networks by non-motor crosslinkers influences cortical mechanics in live cells remains unclear. Moreover, although an “optimal” cortical thickness is required to maintain cortical tension^52^, crosslinkers’ specific contribution to regulating this parameter *in vivo* remains largely unexplored.

To address these questions, we combined optogenetic actin crosslinking with a magnetic pincher to quantify both cortical rheology and thickness in live cells with high spatial and temporal precision^53,54^. To keep a constant cell geometry between our measurements, and facilitate pincher experiments, we plated cells on circular micropatterns of diameter 20µm, close to the diameter of the cell in suspension^55^. Experiments were performed in clonal cell lines with low and high expression of optogenetic crosslinkers, enabling the exploration of the role of crosslinker density in cortical mechanics. We use “density” henceforth to describe conditions of high or low expression. Two experimental constraints prevented magnetic pincher measurements of 7X-CRY2 at high density: the construct could not be stably expressed at high levels long enough to establish clonal lines, and sorting high-expressing cells was incompatible with the magnetic bead incubation required for pincher experiments.

For all experiments, we combined seconds-scale activation (1 pulse/20s) with compression/relaxation cycles (see **Methods**) and measured the same cell before and during activation (**Fig. 2A**). Cortical thickness was quantified prior to each compression (undeformed thickness), while linear fits were applied to force-thickness curves to extract the elastic modulus for individual compressions (**Fig. 2A**). At high density, activation of 1X-CRY2 increases thickness, while 0X-CRY2 had no effect (**Fig. 2B, Fig. S2A**). Measuring cortical stiffness over the 0-500 pN deformation range, we found that 0X-CRY2 induces a small but significant increase, while 1X-CRY2 had no effect (**Fig. S2B, C**). Since stiffness and thickness are correlated (**Fig. S2D**), we normalized the elastic modulus to a common reference thickness (500 nm) to isolate length-specific effects on stiffness (E_norm_). Both 0X-CRY2 and 1X-CRY2 increased E_norm_, with 1X-CRY2 producing a larger increase (**Fig. 2C, Fig. S3E**). This length-dependence in mechanics appears to be density-dependent (**Fig. S3A, B**). Together, these results show that crosslinker length shapes cortical mechanics through distinct routes: shorter crosslinkers primarily increase stiffness, while longer crosslinkers primarily increase thickness.

**Figure 2.**
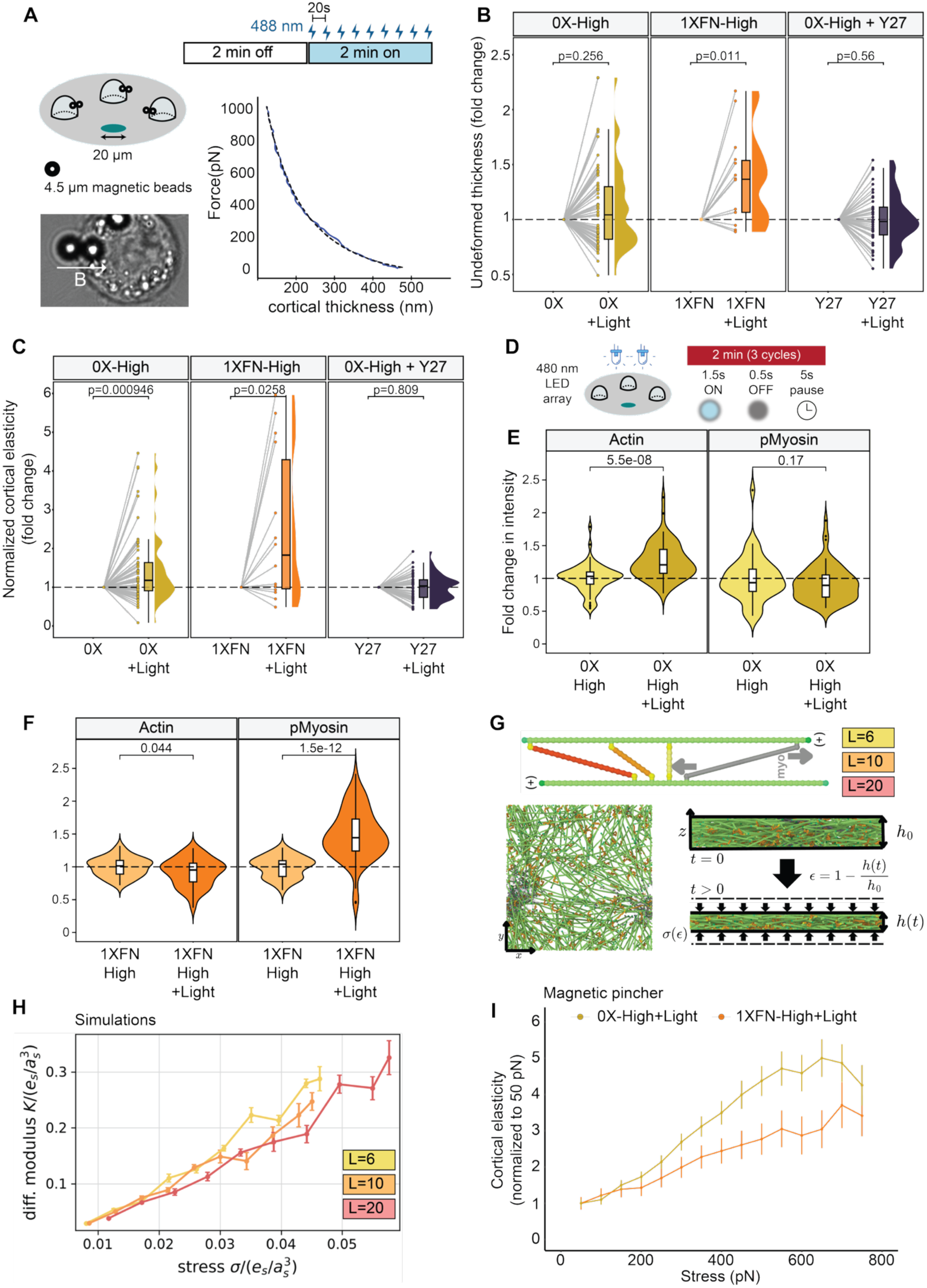
Optogenetic activation of actin crosslinking changes cortical mechanics in a length and myosin-dependent manner. **A)** Schematic of a magnetic pincher experiment. (**left**) NIH-3T3 fibroblasts seeded on 20 µm round micropatterns phagocytose 4.5 µm magnetic beads to pinch the cortex (**bottom**, brightfield image). B depicts the direction of the magnetic field. (**top**) Optogenetic activation protocol for all the pincher experiments. (**right**) Force-distance curve corresponding to one compression. **B, C)** Fold variation of undeformed thickness **(B)** and normalized cortical elastic modulus **(C)** for 0X-CRY2-High (yellow),1XFN-CRY2-High (orange) and 0X-CRY2- High+Y27 (dark blue), normalized to the respective baseline. Each dot represents the average value for one cell. **D)** Schematic of a high-throughput LED array experiment: NIH-3T3 fibroblasts seeded on 20 µm round micropatterns are illuminated over 3 cycles of ON-OFF LED pulses. **E, F)** Relative change in mean cortical intensity for actin and phospho-myosin for 0X-CRY2-High **(E)** and 1XFN-CRY2-High **(F)**, normalized to the control condition (no illumination). Violin plots represent the distribution of all the cells. **G)** MD simulations of cortical compression under linkers of different lengths. **(top)** Actin (green) is modelled as stiff, long filaments with high bending rigidity. Myosin (grey) can bind to actin and in the bound state “walks” along actin-filaments towards the barded (+) end at constant rate inducing active stress. Monodisperse linkers of varying length (6 in yellow, 10 in orange, 20 in red) are modelled as non-rigid polymer chains with two actin binding heads. **(bottom left)** Top-view of a system at steady-state with periodic boundary conditions in x- and y-direction. **(bottom right)** Compression simulations: Top and bottom of the simulation box in z-direction are hard wall boundaries confining the system to an anisotropic configuration. During compression the height of the box is decreased at a constant rate, corresponding to a linear increase in strain. The forces acting on the walls opposing the compaction probe the stress-strain response of the system. **H)** Stress-stiffening curves obtained from simulations, where differential modulus K(\sigma) is derived from stress-strain curves, representing the three different lengths: 6 (yellow), 10 (orange), and 20 (red). **I)** Stress-stiffening curve from magnetic pincher data for 0X-High (yellow) and 1XFN-CRY2-High (orange) during activation, normalized to the differential elastic modulus at 50 pN. Differential elastic modulus was measured at bins of 50pN between 50-700pN. Mean of all compressed cells ± SEM. N_exp_ ≥ 3 independent experiments per condition. P-values are obtained using a paired (Fig. 2B, C) or unpaired (Fig. 2E, F) Wilcoxon-Signed Rank Test. See also Figures S2, S3

**Figure S2.**
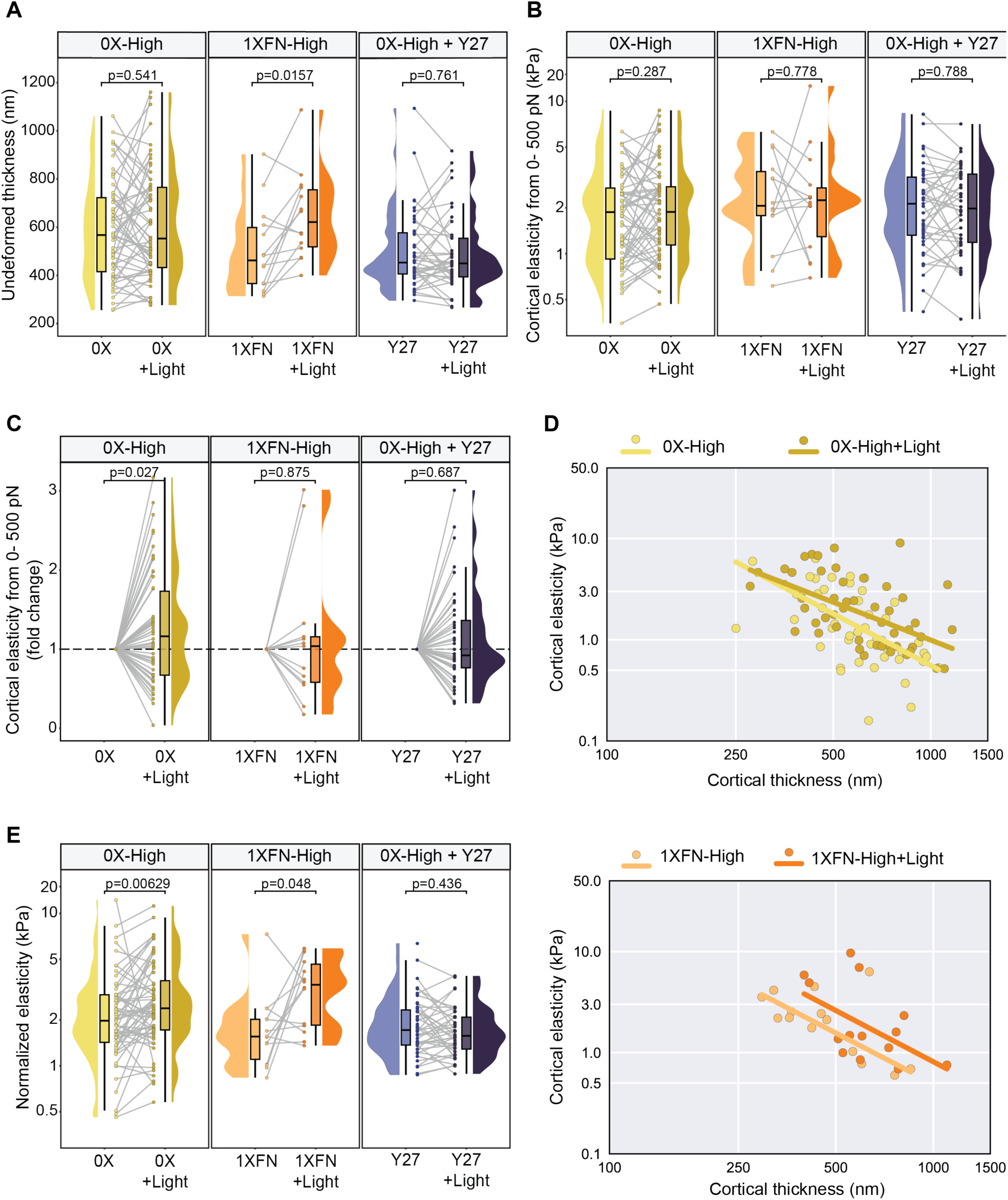
Raw mechanical data and the stiffness-thickness correlation underlying 1XFN’s normalized stiffness effect. **A, B)** Undeformed cortical thickness **(A)** and linear cortical elastic modulus **(B)**, measured between 0-500 pN of deformation, for 0X-CRY2-High (yellow), 1XFN-CRY2-High (orange) and 0X-CRY2-High+Y27 (dark blue). Each dot represents the average value for one cell. **C)** Fold variation of linear cortical elastic modulus (0- 500pN) for 0X-CRY2-High (yellow), 1XFN-CRY2-High (orange) and 0X-CRY2-High+Y27 (dark blue), normalized to the respective baseline. Each dot represents one cell. **D)** Stiffness-thickness correlations for the entire range of measured thickness for 0X-CRY2-High (yellow, before and during activation) and 1XFN-CRY2- High (orange, before and during activation). **E)** Normalized cortical elastic modulus for 0X-CRY2-High (yellow), 1XFN-ECRY2-High (orange) and 0X-CRY2-High+Y27 (dark blue). Y axis plotted as a log_10_ scale for **(B)** and **(E)**. N_exp_ ≥ 3 independent experiments per condition. P-values are obtained using a paired Wilcoxon-Signed Rank Test.

**Figure S3.**
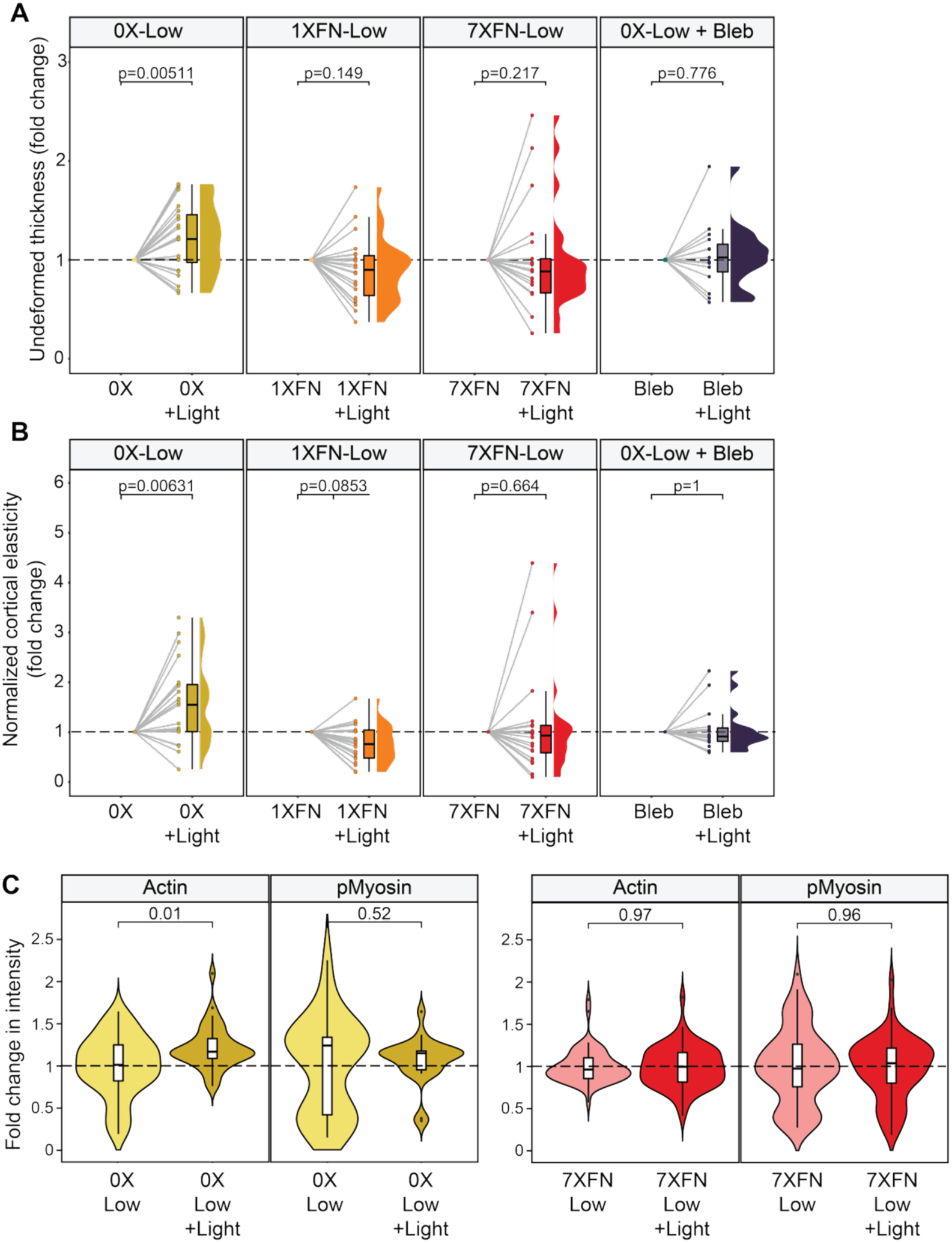
Crosslinker length has a mild impact in cortical mechanics and composition at low density. **A, B)** Fold variation of undeformed thickness **(A)** and normalized cortical elastic modulus **(B)** for 0X-CRY2 low density (yellow), 1XFN-CRY2 low density (orange), 7XFN-CRY2 low density (red) and 0X-CRY2 + Blebbistatin 50 µm (dark blue), normalized to the respective baseline. Each dot represents the average value for one cell.**C)** Relative change in mean cortical intensity for actin and phospho-myosin for 0X-CRY2 low density (left) and 7XFN-CRY2 low density (right), normalized to the control condition (no illumination). Violin plots represent the distribution of all the cells. N_exp_ ≥ 3 independent experiments per condition. P-values are obtained using a paired (Fig. S3A, B) or unpaired (Fig. 3C) Wilcoxon-Signed Rank Test.

Myosin motors are known to stiffen actin networks by pulling on filaments and increasing tension. We therefore asked whether myosin contractility also underlies the crosslinker-driven changes in cortical mechanics we observe. Magnetic pincher experiments on 0X-CRY2 cells treated with the ROCK inhibitor Y-27632 (Y27) showed no stiffness increase driven by light activation, indicating that myosin activity is required for this effect (**Fig. 2B-C**, **Fig. S2B, C, E**). Y27 treatment also reduced undeformed cortical thickness (**Fig. S2A**), suggesting that myosin contractility contributes to baseline cortical thickness independently of crosslinker activation. To more directly test whether crosslinker activation alters myosin or actin recruitment to the cortex, we implemented a LED device to combine high-throughput activation with high-resolution confocal imaging (**Fig. 2D**). We applied a 2 min activation protocol mimicking the pincher activation protocol to micropatterned round cells followed by rapid cell fixation and immunostaining for phospho-myosin (Ser19 site) and actin (**Fig. 2D**). We found that, at high density, short 0X-CRY2 crosslinker promotes cortical actin recruitment (**Fig. 2E**) while mid-length 1X-CRY2 strongly increases phospho-myosin without affecting actin (**Fig. 2F**). Once again, the effect of length was density- dependent (**Fig. S3C**). Therefore, the length-dependent effect of 1X-CRY2 in cortical thickness and E_norm_ (Fig. **2B-C**) is likely associated with higher cortical myosin activity.

Having established that crosslinker length shapes cortical mechanics at the two lengths accessible to our pincher measurements, we next asked whether this relationship holds continuously across the full range of physiological lengths. Together with our inability to test 7X-CRY2 at high density, this motivated us to turn to molecular dynamics simulations. We adapted a simulation framework previously used to model crosslinked actomyosin networks^56^ (**Fig. 2G**). Actin filaments (green) are modelled as stiff, semiflexible polymers with high bending rigidity; myosin motors (grey) bind actin filaments and, once bound, walk toward the barbed (+) end at a constant rate, generating active stress. Crosslinkers of varying length (yellow/orange/red) are modelled as flexible polymer chains with two actin- binding heads. Simulated networks reach a steady state within a box with periodic boundary conditions in the x- and y-directions (**Fig. 2G**). To probe the network’s mechanical response, we performed compression simulations in which the top and bottom of the simulation box act as hard-wall boundaries; decreasing the box height at a constant rate imposes a linear increase in strain, while the resulting force on the confining walls reports the network’s stress-strain response (**Fig. 2G**). This computational approach let us ask a question our experiments alone could not answer: how does stress-stiffening evolve continuously across the full range of crosslinker lengths? Using the differential modulus obtained from stress-strain curves (**Fig. 2H**), we found that the shortest linkers display more non-linear stress-stiffening, while longer linkers exhibit a more linear relationship upon increases of stress. This simulation prediction was directly testable at the two lengths available to us experimentally. We then applied a similar framework to our experimental pincher data, extracting a differential modulus from stress-strain curves, where strain is extracted from changes in thickness (**Fig. 2A, I**). At high density, 0X-CRY2 induces a more nonlinear stress-stiffening response compared to 1X-CRY2, thereby confirming the predictions from our simulations (**Fig. 2H, I**). Together, these results demonstrate that the cortex’s stress-stiffening varies continuously with crosslinker length.

### Crosslinkers trigger cortical actin polarization or delamination in a length-dependent manner

Generation and transmission of myosin-dependent cortical flows is critical to drive cell polarization, and requires constant regulation of actin assembly, crosslinking, density, and filament length^57^. To understand how spatiotemporal regulation of crosslinkers of different lengths contributes to this interplay, we combined optogenetic activation of 0X, 1X, and 7X-CRY2 with fast AiryScan imaging in round micropatterned cells over longer timescales. Our initial seconds-scale illumination protocol delivered too much light, causing the crosslinker to form large clusters; actin turnover then caused actin to be lost from the clusters (**Fig. S4**). Thus, we designed a milder minutes-scale pipeline better suited to the longer timescales needed for cell dynamics. Our illumination protocol comprises an initial, baseline “dark” phase of 4 minutes to account for fluctuations in cortical dynamics, followed by an activation phase with 488 nm light at 1 pulse/min over 6 minutes. This activation window spans multiple cycles of cortical actin turnover in round mammalian cells (15–25 s)^58^. The 1 pulse/min frequency for CRY2 activation was chosen based on previous studies with optogenetic tools and cell dynamics^59^ as well as the kinetics of the tool itself^42,43^.

We found that activation of crosslinkers induced two distinct length-dependent cortical behaviours: cortical actin delamination and cortical actin polarization (**Fig. 3A**). Activation of short 0X-CRY2 and intermediate 1X-CRY2 crosslinkers resulted primarily in cortical actin delamination (**Fig. 3A, B**). Strikingly, activation of long crosslinkers triggered cortical actin polarization in most cells, suggesting that the presence of long crosslinkers is sufficient to drive symmetry breaking (**Fig. 3A, B**). Moreover, both phenotypes were more prevalent in clonal cell lines with higher expression of our tools (**Fig. 3B**), suggesting that crosslinker density is also a regulatory point in cortical dynamics.

**Figure 3.**
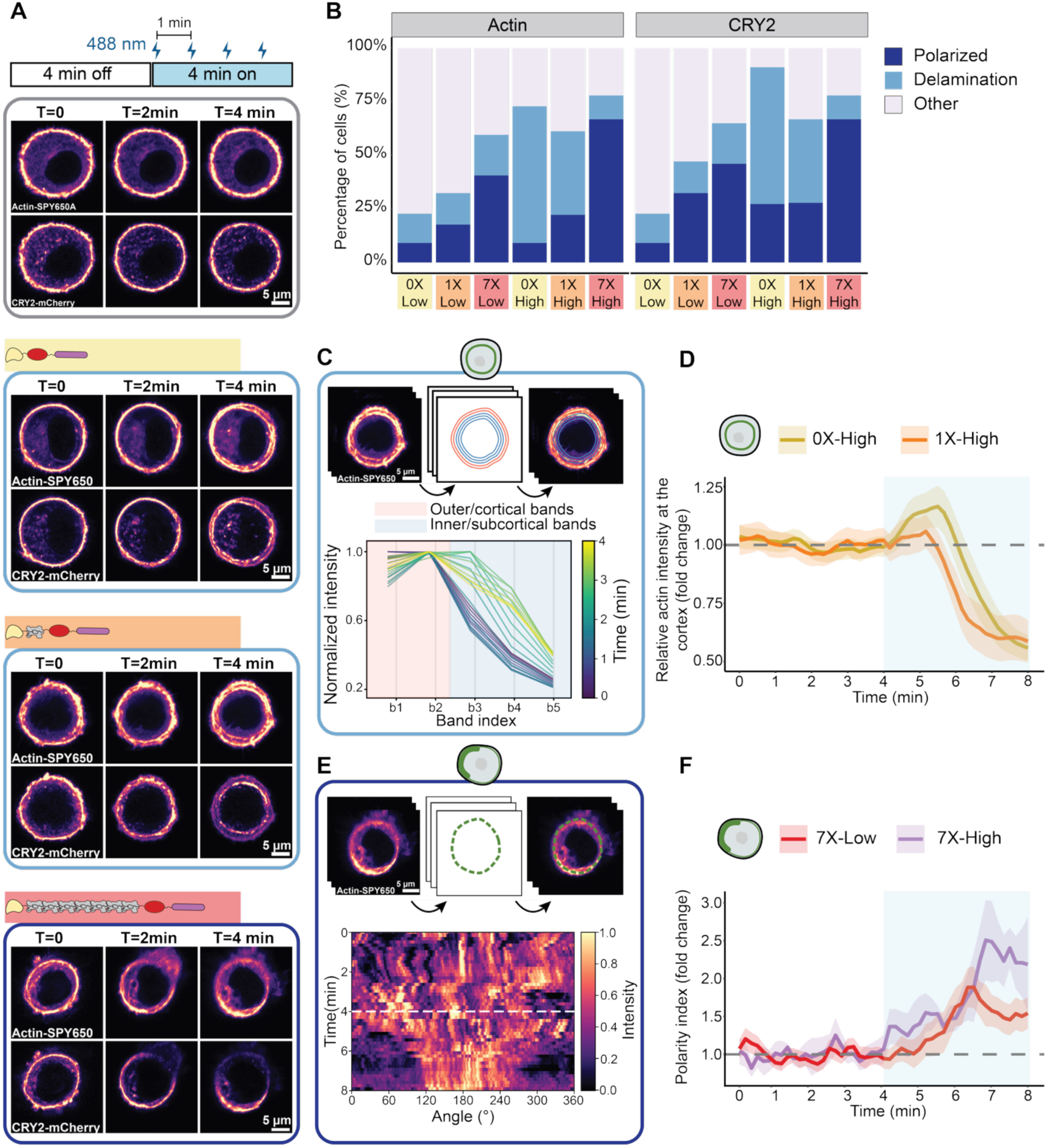
Crosslinker length dictates cortical actin delamination versus polarization, driving symmetry breaking. **A)** Airyscan images of NIH-3T3 fibroblasts expressing 0X-CRY2 (yellow), 1XFN-CRY2 (orange), and 7XFN- CRY2 (red) for high-density conditions after 0, 2, and 4 minutes of activation. For each set of images, top row corresponds to live actin staining with FastAct-SPY650, while bottom row corresponds to mCherry signal of the respective crosslinker. Light blue box highlights delamination for 0X and 1XFN, while dark blue box highlights polarization and symmetry breaking for 7XFN. Light grey box on the top set of images highlights the absence of any of the previous phenotypes. **B)** Fraction of polarization (dark blue), delamination (light blue), and remaining (light grey) phenotypes for all crosslinkers, at low- and high-density conditions, both for actin and CRY2 signal. **C)** Quantification pipeline for delamination. **(top)** Actin-SPY650 channel is segmented, and a series of concentric masks (bands) are traced across all frames. **(bottom)** Normalized intensity across the first five bands. As activation proceeds, the intensity of the sub-cortical bands increase, corresponding to delamination. The ratio between the two most-outer, cortical bands (red) and the subsequent three sub-cortical bands (blue) represents the enrichment of actin at the cortex. **D)** Evolution of cortical actin enrichment throughout a full baseline-activation cycle for 0X-CRY2 (yellow) and 1XFN-CRY2 (orange), normalized to the baseline condition. Each line represents the average of all cells exhibiting delamination. **E)** Quantification pipeline for polarization. **(top)** Actin-SPY650 channel is segmented, and the outline of the mask is overlaid onto the image stack. Successive image contours are unwrapped and stacked to form kymographs. **(bottom)** Representative kymograph for the 7XFN-CRY2 cell in A, displaying the accumulation of actin into a smaller region over time. A polarity index is extracted as a function of the increased intensity and the size of the accumulation region. **F)** Evolution of polarity index throughout a full baseline-activation cycle for 7XFN-CRY2 low-density (red) and high-density (purple), normalized to the baseline condition. Each line represents the average of all cells exhibiting a final polarity index above the cutoff of 0.05 for each condition. See also Figures S4 and S5.

**Figure S4.**
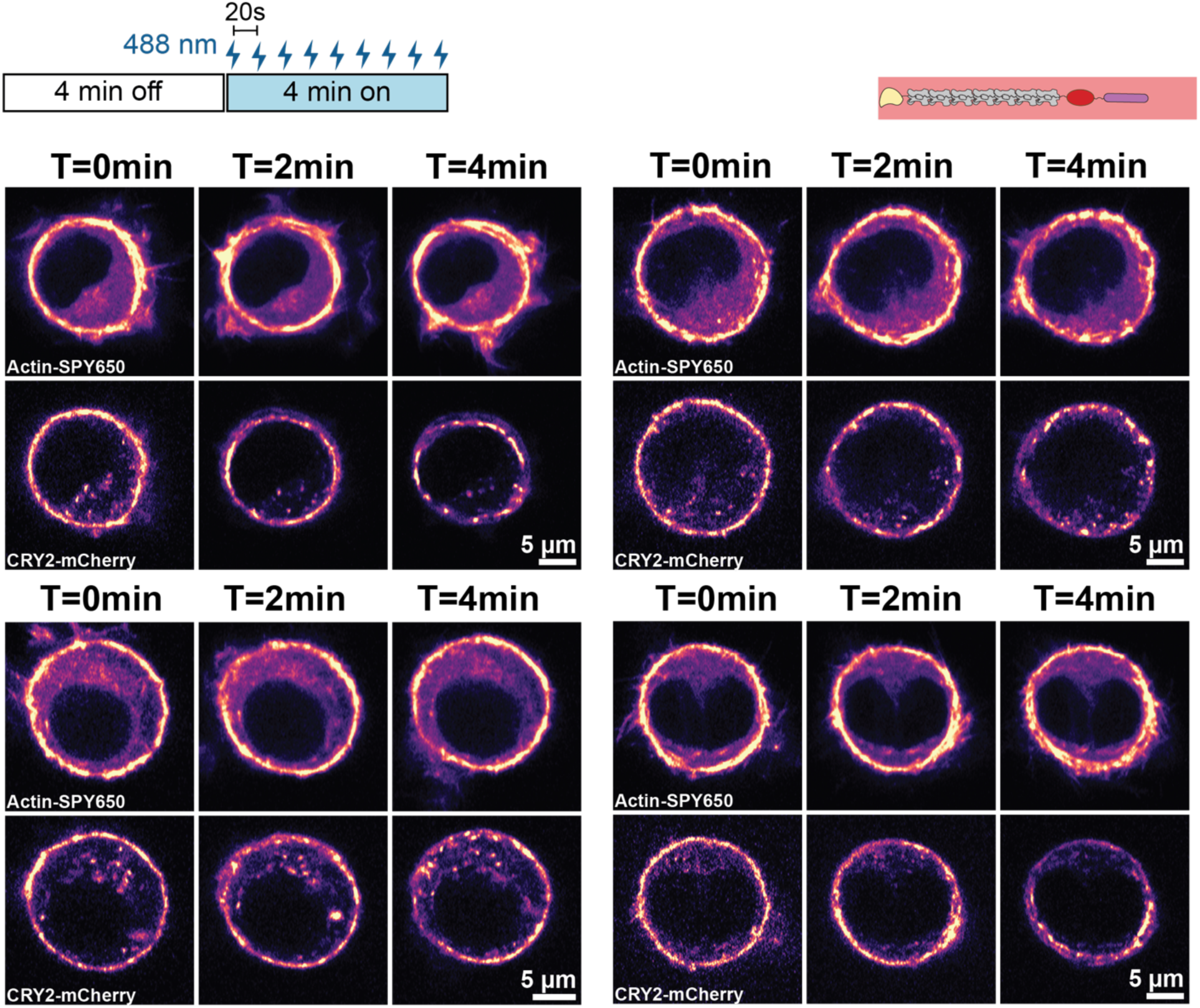
Fast activation leads to clustering and fragmentation of 7X-CRY2 without regulating actin dynamics. Airyscan images of NIH-3T3 fibroblasts expressing 7XFN-CRY2 (red) for low-density condition at 0, 2, and 4 minutes of fast activation (1 pulse/20s). For each set of images, top row corresponds to live actin staining with FastAct-SPY650, while bottom row corresponds to mCherry signal of the 7XFN-CRY2 crosslinker.

**Figure S5.**
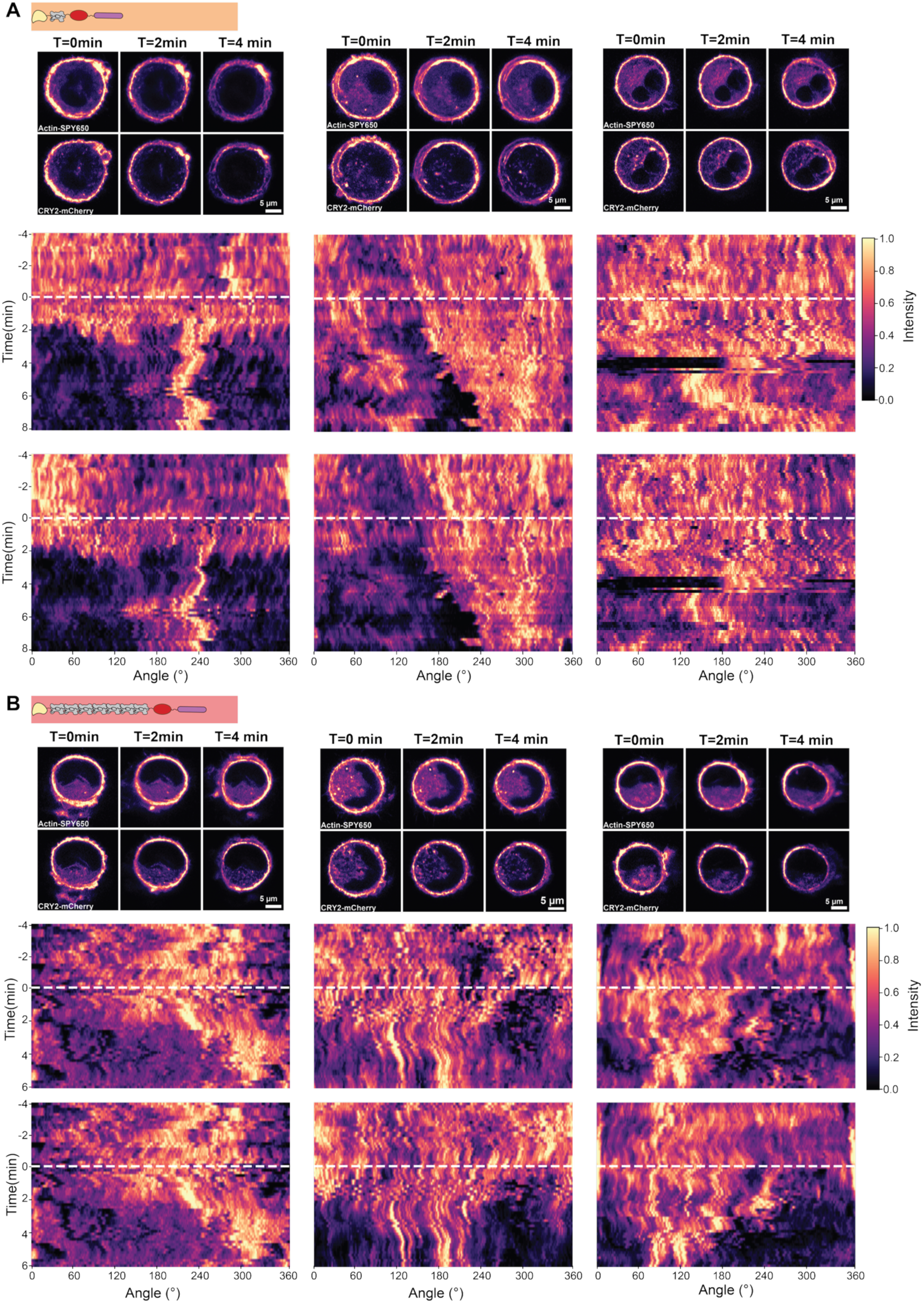
Crosslinker-driven actin polarization exhibits a variety of patterns. **A)** Different polarization patterns for 1XFN-CRY2. (**top**) Airyscan images of NIH-3T3 fibroblasts expressing 1XFN-CRY2 (orange) for high-density conditions at 0, 2, and 4 minutes of activation. For each set of images, top row corresponds to live actin staining with FastAct-SPY650, while bottom row corresponds to mCherry signal of the 1XFN-CRY2 crosslinker. (**bottom**) Correspondent kymographs for each set of cells, obtained from unwrapping and stacking the cell contour for each timelapse point. For each cell, top kymograph corresponds to FastAct-SPY650 while bottom kymograph corresponds to 1XFN-CRY2-mCherry. **B)** Same as A but for 7XFN-CRY2.

Next, we quantified the temporal evolution of both delamination and polarization phenotypes. To quantify delamination over time, we segmented the actin channel at the cortex and traced a series of concentric bands to measure the intensity ratio between the two outer/cortical bands and three subsequent inner/subcortical bands (**Fig. 3C**). 0X-CRY2 triggers an increase of relative cortical actin intensity during the first 2 minutes of activation, followed by a fast decrease marking delamination onset. 1X-CRY2, by contrast, showed a decrease in cortical actin intensity from the start of activation, with delamination onset occurring within the first minute, indicating that the progression of delamination is itself length-dependent. The transient recruitment of cortical actin seen for 0X-CRY2 (**Fig. 3C**) is consistent with the increase in cortical actin observed with 0X-CRY2 activation in our LED-array experiments after 2 minutes of activation (**Fig. 2F**), while 1X-CRY2 failed to elicit a similar effect under either the LED-array protocol (**Fig. 2F**) or the minutes-scale AiryScan protocol used here (**Fig. 3C**).

To measure how polarization evolves over time, we segmented the actin channel and dilated the segmented mask to include the cortex. We then unwrapped this band and stacked it over time to obtain kymographs (**Fig. 3E**), which are divided into angular bins (see **Methods).** Next, we extract both the local enrichment of the signal at the cortex and the degree of angular enrichment, which are used to compute a polarization index, reflecting both the intensity and spatial distribution of actin enrichment at the cortex (see **Methods**) (**Fig. 3F**). Quantification of the polarization index over time revealed a progressive increase of cortical actin polarization throughout the first 2-3 minutes of activation followed by a plateau. Moreover, high-expression 7X-CRY2 cells displayed a larger increase in polarization index with an earlier onset, consistent with a higher fraction of polarized cells compared to low expression conditions. Visual inspection of kymographs for polarized 1X-CRY2 (**Fig. S5A**) or 7X-CRY2 (**Fig. S5B**) cells showed that the region where polarization emerges is not fixed in advance, but is instead established *de novo* upon activation, rather than reinforcing pre-existing patterns of actin accumulation. Altogether, our results show that crosslinker length dictates whether the cortex is remodelled to promote delamination or polarization, with long crosslinkers being sufficient to induce symmetry breaking.

### Cortical actin polarization and delamination depend on actomyosin contractility and actin polymerization

Our live imaging data shows that long 7X crosslinkers can drive cortical actin polarization and symmetry breaking, while shorter 0X and 1X crosslinkers primarily lead to delamination. As both outcomes are forms of cortical flow, and these are generated by actomyosin contractility, we tested whether myosin activity regulates both phenotypes. To that end, we combined crosslinker activation and live imaging in the presence of the myosin inhibitor Blebbistatin (20 µM). Treatment with blebbistatin completely prevented delamination and polarization for 1X-CRY2, with cortical actin remaining stable throughout the entire activation period (**Fig. 4A, B, Fig. S6**). Conversely, blebbistatin only slightly reduced the frequency of delamination for 0X-CRY2 cells (**Fig. 4A, B, Fig. S4A**). In cells that still delaminated, onset was delayed by ∼1 minute relative to control, though it was still preceded by the initial increase in cortical actin recruitment seen in untreated cells. Last, when combined with 7X-CRY2 activation, blebbistatin treatment strongly reduced the fraction of polarized cells as well as the polarization index, which remained low throughout the activation period (**Fig. 4A, B, Fig. S6**). Together, these results confirm that polarization depends on myosin activity, while myosin’s contribution to delamination varies with crosslinker length.

**Figure 4.**
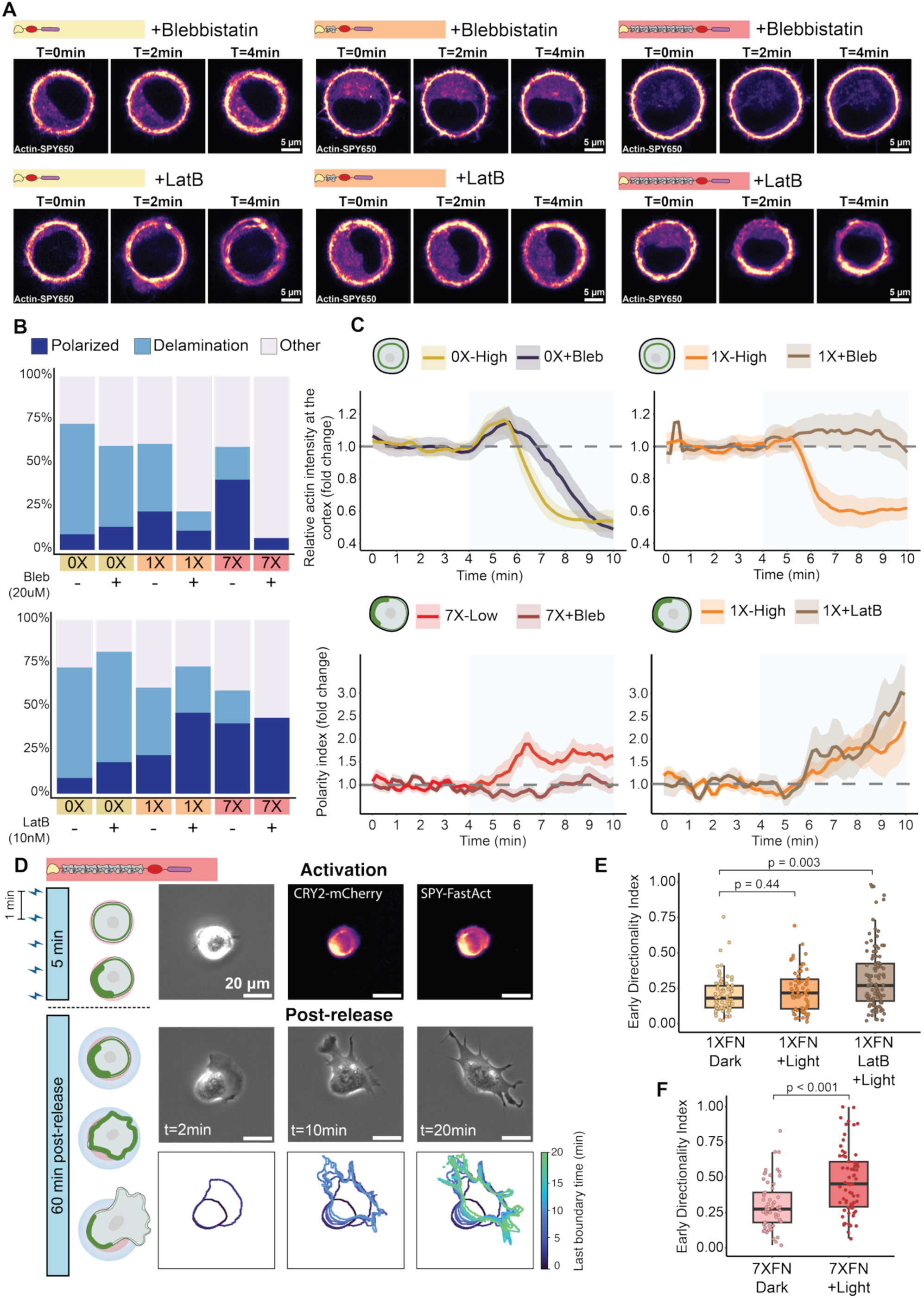
Polarization is dependent on myosin activity and actin connectivity, ultimately directing cell motility. **A)** Airyscan images of NIH-3T3 fibroblasts with live actin FastAct-SPY650 staining expressing 0X-CRY2 (yellow), 1XFN-CRY2 (orange), and 7XFN-CRY2 (red) at 0, 2, and 4 minutes of activation in the presence of the myosin inhibitor Blebbistatin (20 µM) (**top**) or the inhibitor of actin polymerization Latrunculin B (10 nM) (**bottom**). **B**) Fraction of polarization (dark blue), delamination (light blue), and remaining (light grey) phenotypes for all crosslinkers with 20 µM Blebbistatin (**top**) or 10 nM Latrunculin B (**bottom**). **C**) (**top**) Evolution of cortical actin enrichment throughout a full baseline-activation cycle for 0X-CRY2 and 1XFN-CRY2, normalized to the baseline condition, in the presence of 20 µM Blebbistatin (yellow for 0X-CRY2, dark blue for 0X-CRY2 with Blebbistatin, orange for 1XFN-CRY2, light brown for 1XFN-CRY2 with Blebbistatin). Each line represents the average of all cells exhibiting delamination for control conditions and a randomly selected set of cells for drug treated conditions. (**bottom**) Evolution of polarity index throughout a full baseline-activation cycle for 7X-CRY2 and 1XFN-CRY2, normalized to the baseline condition in the presence of 20 µM Blebbistatin (red for 7XFN-CRY2, dark brown for 7XFN-CRY2 with blebbistatin) or 10 nM Latrunculin B (orange for 1XFN- CRY2, light brown for 1XFN-CRY2 with Latrunculin B). Except for 7XFN-CRY2 with blebbistatin, each line represents the average of all cells exhibiting polarization; for 7XFN-CRY2 with blebbistatin, lines correspond to randomly selected cells. **D**) Dynamic micropatterning experiments to characterize cell spreading and motility. (**top**) Brightfield and epifluorescence images of an NIH-3T3 fibroblast expressing 7XFN-CRY2 after a 5 min activation cycle with 1 pulse/min. (**middle**) Brightfield images of the same cell at 2, 10, and 20 min after pattern release with SA-Fibronectin. (**bottom**) Respective cell contours overlaid and color-coded across time. **E**) Directionality index calculated for the first 10 minutes after pattern release for 1XFN-CRY2 at control/dark (light orange), activation (orange) and Latrunculin B-treated (light brown) conditions. **F**) Directionality index calculated for the first 10 minutes after pattern release for 7XFN-CRY2 at control/dark (light red) and activation (dark red) conditions. Each dot represents one cell. There are 3 independent experiments per condition. P-values are obtained using a Linear Mixed Model. See also Figure S6.

**Figure S6.**
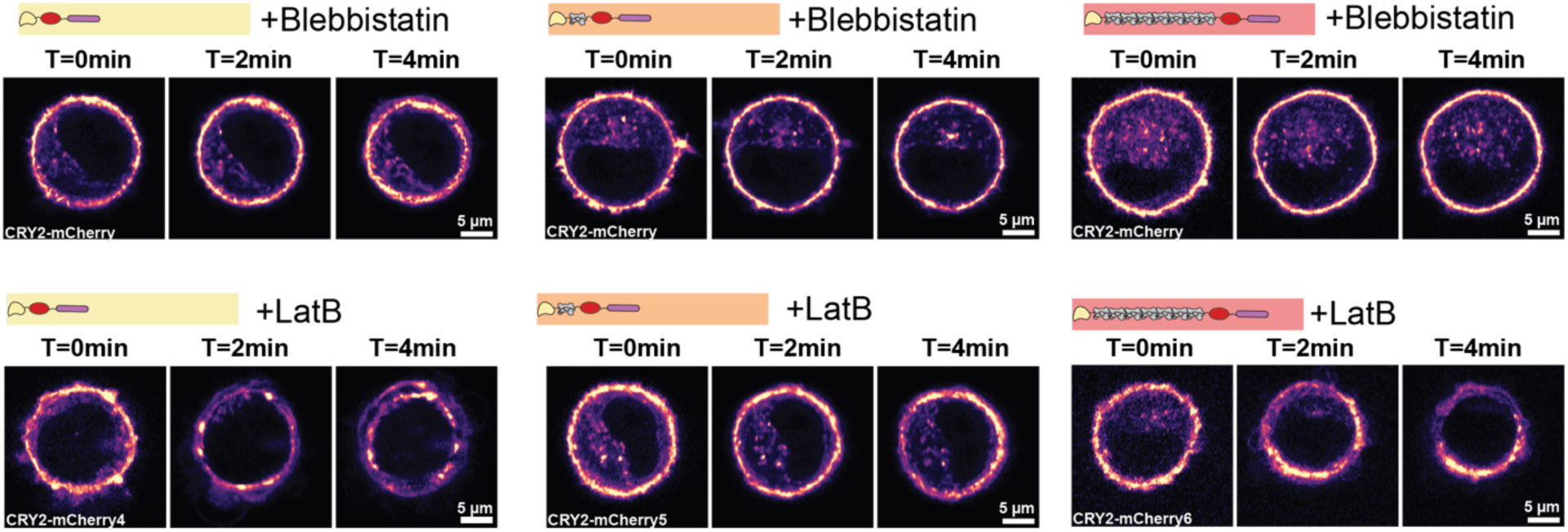
Polarization is dependent on myosin activity and actin connectivity. Airyscan images of NIH-3T3 fibroblasts expressing 0X-CRY2 (yellow), 1XFN-CRY2 (orange), and 7XFN-CRY2 (red) at 0, 2, and 4 minutes of activation in the presence of myosin inhibitor Blebbistatin (20 µM) (**top**) or inhibitor of actin polymerization Latrunculin B (10 nM) (**bottom**).

Actin network connectivity is further modulated by turnover of both crosslinkers and actin filaments, which in turn can stabilize contractile steady states^6^, promote cyclic/pulsed contractions^60^, and change cortical flows^61^. Disruption of actin polymerization or depolymerization both lead to changes in actin turnover^58,61^, which led us to assess the effect of optogenetic crosslinkers under perturbations of actin polymerization. To this end, we used low dose of Latrunculin B (10 nM), a drug which binds G-actin monomers, preventing their incorporation into actin filaments; a low dose ensures that actin polymerization is mildly affected while retaining the global network. Treatment with 10 nM LatB did not substantially alter the behaviour for 0X-CRY2 and 7X-CRY2, with delamination and polarization still being the main phenotypes, respectively. However, Latrunculin B treatment doubled the fraction of polarized cells for 1X-CRY2 while reducing the fraction of cells undergoing delamination (**Fig. 4A, B**). The onset and progression of polarization followed a similar pattern as the control condition while the polarization index was higher, suggesting a stronger local accumulation of actin. Therefore, perturbations of actin polymerization can promote crosslinker-induced symmetry breaking but only for intermediate-length crosslinkers.

### Length-driven actin polarization directs cell motility

Establishment and maintenance of cell polarity is a key step in driving persistent, directed cell motility. In mesenchymal cell migration, front-to-rear polarity can be established by a combination of mechanisms including biochemical gradients of Rho GTPases^62^, membrane tension propagation^63,64^, and contractility^64^, among others. However, it is unclear whether cortical actin polarization induced by long crosslinkers could also contribute to directed cell motility. We tested this hypothesis by combining optogenetic activation of long crosslinkers with dynamic micropatterning based on ultraviolet (UV) photopatterning of biotinylated PLL-g-PEG^65^. By using streptavidin-conjugated secondary ligands such as fibronectin, this method allows conventional micropatterning of adhesive and non-adhesive regions coupled to selective control of cell adhesion over time. By timing fibronectin addition relative to optogenetic activation, cells can be released from spatial confinement to adhere and spread onto a new ligand while crosslinker activity is being manipulated (**Fig. 4D**).

Cells expressing 7X-CRY2 were plated in round micropatterns and activated over 4 minutes (**Fig. 4D**) to match the maximum increase of polarity index measured by live imaging (**Fig. 3D**). We then quantified cell spreading during the subsequent release from the micropattern. Quantification of spreading in the 10 minutes post-release revealed that activation of 7X-CRY2 crosslinkers leads to a significant increase in spreading directionality compared to control (**Fig. 4D, F**), confirming that polarization driven by long crosslinkers directs cell motility.

Next, we investigated whether LatB-driven polarization for 1X-CRY2 is also sufficient to regulate cell spreading and motility. Dynamic micropatterning experiments showed that 1X-CRY2 cells display a mild but non-significant increase in spreading directionality upon activation compared to the dark condition (**Fig. 4E**). Notably, 1X-CRY2 cells co-treated with LatB exhibited a significantly higher spreading directionality upon activation, confirming that LatB-induced polarization in 1X-CRY2 cells also directs cell motility (**Fig. 4E**). Overall, our experiments show that long and mid-length crosslinkers can dynamically remodel cortical actin to promote symmetry breaking and ultimately establish cell polarity, resulting in directed cell motility.

## DISCUSSION

Our results show that crosslinker length, on its own, is sufficient to switch the cortex between distinct mechanical and morphogenetic states, from reinforcement to collapse to polarization, ultimately directing cell motility. At second timescales, short and mid-length crosslinkers reinforce the cortex, increasing its stiffness and thickness respectively, with shorter crosslinkers driving more pronounced stress-stiffening, a relationship simulations show holds continuously across the full range of crosslinker lengths. These effects require myosin contractility and are accompanied by crosslinker-driven changes in cortical composition. Strikingly, at minute timescales, crosslinker length no longer simply reinforces the cortex, but instead determines whether it collapses or polarizes; short crosslinkers drive cortical delamination, while long crosslinkers trigger polarization and symmetry breaking. Myosin contractility remains essential to this switch, as its inhibition abolishes symmetry breaking altogether. Perturbing actin polymerization, however, can override crosslinker identity: mid-length crosslinkers, which normally delaminate, instead adopt the polarizing behaviour otherwise reserved for the longest crosslinker. Together, these results show that crosslinker-induced symmetry breaking, whether set by length alone or unlocked by perturbed actin polymerization, is sufficient to direct cell motility. Myosin contractility emerges as a common thread throughout, required for cortical mechanics, delamination, and polarization alike — but it is the material architecture set by crosslinker length that determines how, when, and where this contractile machinery gets deployed. Length offers cells a fundamentally different kind of control than concentration: a structurally fixed, molecularly encoded switch, rather than one requiring active, ongoing regulation of crosslinker abundance.

### Crosslinker length tunes cortical mechanics across scales

The idea that crosslinkers shape network mechanics is almost as old as their discovery. Already in 1965^66^, when alpha-actinin was first isolated, its authors proposed that it could influence actin network contraction and viscosity^66,67^. Over the course of many subsequent decades, *in vitro* studies have established how density, length and flexibility of actin crosslinkers can regulate actin network strain-stiffening, force transmission, and hysteresis^16,29,33,68–70^. However, studies on the impact of cortical actin structure in live cell mechanics have focused instead on actin filament length^52,71^, while crosslinkers’ contribution has received far less attention. Existing evidence for a crosslinker-specific role is, moreover, limited and difficult to compare directly: alpha-actinin for example accumulates in response to cortical deformation^72^, and its depletion reduces cortical tension in *Dictyostelium*^73^, yet its depletion has no detectable effect on cortical thickness in mitotic HeLa cells^52^. Together, these fragmented observations point to a deeper problem: no system has yet allowed crosslinker length and density to be manipulated with precision in live cells, while directly resolving the resulting mechanical consequences. Our optogenetic toolbox, coupled with magnetic pincher measurements, provides exactly this capability: it allows crosslinker length to be tuned precisely while directly measuring its mechanical impact at second timescales in live cells. Using this approach, we find that crosslinker length shapes both the material (stiffness) and physical (thickness) properties of the cortex. These results demonstrate that the molecular organization of cortical components (here set by crosslinker length and the resulting filament connectivity and network architecture) directly determines the global mechanical properties of the cell.

### Thickness and stiffness arise through distinct, myosin-dependent mechanisms

While crosslinker length clearly shapes the cortex’s mechanical properties, the molecular routes behind this are likely distinct for each parameter. We find that 1X-CRY2, but not 0X-CRY2, increases cortical thickness (**Fig. 2B**). Changes in thickness could reflect the cortical phospho-myosin recruitment we observe specifically for 1X-CRY2 (**Fig. 2F**), since myosin II inhibition is known to reduce both the magnitude and fluctuations of cortical thickness^74^. Notably, fimbrin, which is similar in length to 1X-CRY2, has been shown to recruit myosin II to high-tension cortical domains at epithelial junctions^10^. This raises the possibility that crosslinkers of this specific length are particularly well-suited to promote myosin recruitment through the network architecture they generate^75^.

Myosin contractility also appears to underlie the observed increase in cortical stiffness: myosin motors stiffen actin networks by pulling on filaments^76^, and consistent with this, the ROCK inhibitor Y27 abolishes the stiffness increase driven by 0X-CRY2 (**Fig. 2C**). Additionally, the recruitment of cortical actin we observe for 0X-CRY2, in both high-throughput (**Fig. 2E**) and live imaging (**Fig. 3D**) experiments, could independently contribute to its increase in network stiffness. This would be in line with previous work showing that increased cortical actin, driven by depletion of barbed-end capping and actin-severing proteins, similarly increases cortical thickness^52,77^. By the same logic, the increase in cortical actin we observe for 0X-CRY2 could reflect reduced actin depolymerization, rather than increased polymerization, at the cortex.

### Cortical stress-stiffening is a continuously tuneable, length-dependent property

Beyond setting the magnitude of cortical stiffness and thickness, crosslinker length also shapes the nonlinearity of the cortex’s mechanical response. However, our discrete constructs alone cannot reveal whether this relationship is smooth or idiosyncratic. Molecular dynamics simulations, parameterized to match our experimental system, resolved this: modelling crosslinkers across a broader range of lengths shows that stress-stiffening declines steadily with increasing length (**Fig. 2G, H**). This trend is confirmed experimentally, where 0X-CRY2 shows a more non-linear response than 1X-CRY2 (**Fig. 2I**). This length-dependent stiffening is consistent with established polymer network theory^78^, and has been observed experimentally in purified actin networks with synthetic crosslinkers of varying length^70^. Our results show that this same physical relationship holds in the living cell cortex, despite its far greater molecular complexity and continuous turnover.

In other cytoskeletal systems, nonlinear stiffening under deformation is thought to act as a protective mechanical fuse: vimentin networks stiffen to shield the nucleus from deformation^79^. However, this protection is not always permanent: sustained mechanical loading can trigger structural transitions that undo it, as recently shown for keratin networks, which reorganize and release their protective grip on the nucleus under prolonged stretch^80^. Our toolbox is well placed to test whether short crosslinkers’ pronounced stress-stiffening reflects a protective response, and whether its failure under sustained load explains delamination.

### A length-dependent switch drives cortical symmetry breaking

Over longer, minute-scale activation, crosslinker length governs whether the cortex remains intact or breaks symmetry entirely. Almost 40 years ago, Bray and White proposed that cortical actin flows arise from tension gradients across the cortex, which draw cortical material from relaxed regions toward regions under contraction^81^. These flows (driven by tension anisotropy^82^, turnover^61,83^, and connectivity^57,84^) are critical for symmetry breaking and cell polarization during processes such as division and motility^82,85^. Actin crosslinkers are strong candidates to regulate these flows, since they directly control network connectivity. Indeed, adding alpha-actinin can trigger symmetry breaking in reconstituted contractile cortices^86^. Whether this holds in live cells, however, has been difficult to test directly — until now.

In our experimental setting, crosslinker length alone was sufficient to dictate whether cells polarized or lost cortical integrity: long 7X-CRY2 crosslinkers drove cortical polarization and symmetry breaking, while short 0X and mid-length 1X-CRY2 crosslinkers instead caused cortical delamination. This length-dependent control of symmetry breaking is consistent with a broader role for molecular length as a regulator of cortical mechanics^56^. We hypothesize that 7X-CRY2 remodels the network to generate localized filament entanglement in a specific region of the cortex, producing a heterogeneous increase in connectivity and a corresponding accumulation of contractility in that region. Since 7X-CRY2 falls in a broadly similar length range to alpha-actinin, the threshold for entanglement and polarization may be set around this general length scale. Delamination, by contrast, may result from more uniform connectivity driven by shorter crosslinkers. Here, contractility instead drives uniform retraction of a portion of the cortex, a phenotype reminiscent of apical fiber collapse at tricellular junctions in *Drosophila*, where localized fibers similarly retract under contractile stress^87^. Such delamination-like phenotypes have not, however, been thoroughly explored in the cortex, and could have broader implications for cell morphology and mechanics.

### Myosin activity and actin turnover set the switch for symmetry breaking

What determines this length-dependent switch between delamination and polarization? Cortical tension, generated through myosin II filament pulling and actin polymerization, has been shown to drive symmetry breaking and polarization^56,82,88–90^. Consistent with this, blebbistatin abolishes 7X-CRY2-driven polarization. Staining for phosphorylated myosin in polarized cells would help clarify whether activated myosin is specifically enriched in the polarized region. Myosin also shapes the delamination phenotype, but in a length-dependent manner: differences in network stiffness between 0X-CRY2 and 1X-CRY2 may explain why myosin II inhibition abolishes delamination in 1X-CRY2 but has a smaller effect on 0X-CRY2. These opposite outcomes could also reflect the differential recruitment of cortical actin versus phospho-myosin driven by 0X- and 1X-CRY2, respectively.

Beyond myosin, actin turnover offers a second lever for tuning 1X-CRY2’s behaviour. Filament turnover is known to be required for active stress generation^83,91^ and cortical remodelling^91^, and low doses of Latrunculin A can trigger and enhance periodic cortical waves in CHO cells^92^. We propose that the length-specific enhancement of polarization we observe for 1X-CRY2 upon mild Latrunculin B treatment arises from a similar interplay between perturbed filament turnover and network connectivity, one that becomes accessible specifically at this intermediate crosslinker length. Distinct barbed-end subpopulations with different turnover kinetics^58^ could further tune this effect, offering a specific target for future work dissecting how turnover and crosslinking jointly drive symmetry breaking.

### Crosslinker-induced symmetry breaking directs cell motility

Beyond its mechanical consequences, this length-dependent symmetry breaking directs cell spreading. Actin cortex organization is central to directed migration: a contractile, crosslinked meshwork at the back squeezes the cell forward^64^. Crosslinkers have been linked to migration, notably filamin, through uropod formation during neutrophil migration^93^, but their role in establishing front-back organization from an unpolarized state was unclear. We show that long crosslinkers alone are sufficient to drive this initial transition, from an unpolarized, adherent cell to a polarized, motile one. Our toolbox now offers a way to dissect how this front-rear architecture first emerges, and how it is subsequently sustained during persistent migration.

We propose that crosslinker-driven polarization generates a contractile “rear” that mechanically propels the cell forward. This is consistent with a broader principle: asymmetries in myosin contractility alone can switch cells between non-motile and motile states, as shown in both *Drosophila*^94^ and epithelial cells^95^, and retrograde actin flow coupled to myosin contraction is known to sustain directed migration^85,96^. We hypothesize that long crosslinkers similarly generate a contractility gradient that stabilizes motility during early spreading, potentially reinforced by biochemical feedback: a recent study showed that the actin cortex and membrane tension jointly pattern Rho/Rac1 activity to coordinate front-back polarity^97^, raising the possibility that crosslinker-driven polarization engages a similar mechanism.

Together, our results establish crosslinker length as a molecular dial that dictates cortical mechanics, symmetry breaking, and cell motility. Nature appears to have mastered a principle that materials science is only beginning to grasp: that a single tunable parameter, adjusted across families of related proteins, can generate a whole spectrum of adaptive material behaviours — steering the cortex through reinforcement, collapse, and polarization, and ultimately directing where a cell moves. As optogenetic and computational tools continue to bridge molecular design and cell behaviour, we anticipate that this length-encoded logic will extend well beyond the cortex, to any cytoskeletal system where architecture itself can serve as a decision-maker.

### Limitations of the study

Several limitations of our study should be noted. First, our optogenetic crosslinkers are synthetic constructs designed to match the length of specific endogenous crosslinkers, though they may not fully recapitulate native flexibility and mechanical properties. Second, our mechanical observations must be interpreted within a broader multiparametric space, since a given mechanical outcome (stiffness, thickness) can arise through multiple, non-exclusive molecular routes, including changes in filament length, crosslinking, or myosin activity; for instance, cortical thickness and mesh size are not always coupled^77^, nor does thickness always track with tension^77^. Although we observe crosslinker length-dependent changes in cortical thickness, we did not directly measure mesh size or filament organization. Therefore, combining our mechanical measurements with structural approaches such as scanning electron microscopy or cryo-electron tomography will be needed to establish whether these thickness changes reflect underlying changes in network architecture. Third, our dynamic micropatterning experiments probe motility in a distinct paradigm from previous studies, which have primarily examined suspended or spread migrating cells; direct comparisons to these systems should therefore be made with caution. Finally, several of the mechanisms we propose to explain our observations (including length-dependent filament entanglement, differential myosin recruitment, and crosslinker-driven Rho/Rac1 spatial patterning) are consistent with our data but were not directly tested here. Our toolbox’s precise temporal control also leaves open whether crosslinker length sets the duration, and not only the magnitude, of a given mechanical response, a role recently proposed for other cytoskeletal networks in timing mechanical signal transmission to the nucleus. Ongoing theoretical and experimental work should help establish which of these mechanisms best account for our observations.

## AUTHOR CONTRIBUTION

F.N.V and A.D-M conceived the project. F.N.V, M.P, O.D.R, J.H, and A.D-M designed the experiments. F.N.V., A.J., and B.V performed the magnetic pincher experiments under the supervision of M.P, O.D.R, and J.H. F.N.V. performed all remaining *in cellulo* experiments. L.F developed pipelines to obtain high expression single cell clones. A.D developed the cloning strategy for the generation of the optogenetic crosslinker and performed RT-DC experiments together with M. K. under the supervision of J.G. R.T.M developed the cloning strategy for the generation of Fibcons of different lengths. I.P., M.W. and M.R. developed the computational models. M.W and M.R developed the computational models and performed and analyzed computer simulations with crosslinkers of various lengths under the supervision of I.P and A.Š. F.N.V and A.D.M wrote the manuscript. All authors contributed to the interpretation of the data, read, edited and approved the final manuscript.

## ACKNOWLEDGMENTS

We thank the Diz-Muñoz lab members for useful discussions. We thank the EMBL Flow Cytometry Core Facility, the Electronics Workshop, and the EMBL Advanced Light Microscopy Facility for support and advice. We thank Alejandro Gil from the Electronics Workshop for manufacturing the LED array, and Aliaksandr Halavatyi and Manuel Gunkel from the EMBL Advanced Light Microscopy Factility for assistance with high-throughput and adaptive feedback microscopy.

Funding: We acknowledge the financial support of the European Molecular Biology Laboratory (EMBL), the Human Frontiers Science Program (HFSP) grant RGY0073/2018, the Deutsche Forschungsgemeinschaft (DFG) SFB 1638/1 grant, the Ben Barres Acceleration award from CZI and the ERC grant 101124221 (MitoMeChAnics) to A.D.-M.. F. N. V. acknowledges postdoctoral fellowships from the European Molecular Biology Organization (EMBO postdoctoral fellowship No. 425-2021) and the Joachim Herz Stiftung (Add-on Fellowship for Interdisciplinary Life Science No. 1850035). A.Š. acknowledges funding from the European Research Council under the European Union’s Horizon 2020 research and innovation programme (ERC grant No. 802960), a Vallee Scholar Award from the Vallee Foundation, as well as support from ISTA. I.P. received support from a Theory@EMBL fellowship.

*This work is partially funded by the European Union. Views and opinions expressed are however those of the author(s) only and do not necessarily reflect those of the European Union or the European Research Council Executive Agency. Neither the European Union nor the granting authority can be held responsible for them*.

## MATERIALS AND METHODS

### Cloning

Utrophin CH1 and CH2 domains (UtCH1&2) and mCherry were PCR amplified from mCherry-UtrCH, a gift from William Bement (Addgene, # 26740). UtCH1&2 and mCherry were restriction cloned into pOVC1, a gift from Harald Janovjak (Addgene, # 131171) to generate UtCH-mCherry or mCherry-UtCH for RT-DC experiments. To generate UtCH-mCherry-UtCH, an additional domain was added via restriction cloning. CRY2 domain was obtained from an optogenetic vector library developed by Harald Janovjak’s Lab ^98^ and restriction cloned into UtCH-mCherry to generate UtCH-mCherry-CRY2.

Fibcon sequences were purchased from GeneScript as pUC57-1XFN. Since Fibcons cannot be PCR amplified, 7XFN was generated by successive restriction cloning of Fibcon sequences following a strategy that allows for iterative chaining of individual repetitive domains^99^. With a combination of enzymes (FauI, AcII and BamHI), we generated sticky compatible ends that after ligation allow for an in-frame insertion with a seamless scar. We used this strategy to obtain pUC57-6FN (combination from 2x pUC57-3XFN) were ligated. The 6FN were then re-ligated into the pUC57-Lyn-1XFN, leaving a puC57- Lyn-7FN domain with no ligation scars for subsequent cloning. Then, UtCH-CRY2-mCherry was edited by site-directed mutagenesis to remove an additional NotI site and then restriction cloned into pUC57 vector. 1X or 7X Fibcons were then inserted between PCR- amplified UtCH and mCherry-CRY2 using Gibson assembly to generate UtCH-1XFN-mCherry-CRY2 and 7XFN-mCherry-CRY2.

### Cell culture

NIH-3T3 fibroblasts (CRL-1658, ATCC repository) were cultured in Dulbecco’s Modified Eagle Medium 292 (DMEM) with 4.5 g/l D-glucose (Merck), supplemented with 10% fetal bovine serum and 1% Penicillin Streptomycin (10,000 U/ml). Cells were cultured at 37 ° C with 5% CO2 on polystyrene 100 mm dishes (Corning). Cells were passaged every 2-3 days (up to 70% confluency) using 0.05% Trypsin-EDTA. Cells were routinely tested for mycoplasma contamination using MycoAlert detection kit (Lonza) according to manufacturer’s instruction.

### Generation of transgenic cell lines

For cell line generation with the PiggyBac transposon system, the gene of interest was cloned via Gibson assembly into a custom PiggyBac vector downstream of a doxycycline (dox)-responsive TRE3G promoter. The transposable region of the vector also contained a TET3G and a Neomycin resistance cassette under the control of a CAG promoter. Stable cell lines were obtained by co-transfecting the PiggyBac vector and the PiggyBac transposase using Lipofectamine 3000 (#L3000-008, Invitrogen). Successfully transfected cells were then positively selected by adding 1.2 mg/mL Geneticin (#10131-035, Gibco) to the medium.

For lentiviral transduction, the gene of interest was cloned by restriction cloning into a pRRL transfer plasmid (LT3GEPIR, #111177, Addgene) downstream of a doxycycline (dox)-responsive TRE3G promoter. The transfer plasmid also contained a Puryomycin-resistance cassette. Lentiviruses were then produced in HEK-293T cells. Cells at ∼60% confluency were transfected with 1.5 μg of pRRL transfer plasmid (containing gene of interest), 0.176 μg of the viral envelope plasmid (pMD2.G, Addgene, #12259) and 1.3 μg of the viral packaging plasmid (pCMV 8.74, Addgene, #22036). Transfection was performed using the TransIT-Lenti reagent (#MIR 6603, Mirus) together with OptiMEM (#31985070, Gibco) according to the manufacturer’s protocol. Infectious viral particles were harvested on day two post-transfection and concentrated 25-30x by using Lenti-X concentrator (#631231, Takara). In brief, the supernatant of transfected HEK-293T cells was filtered through a 0.22 µm PVDF filter and mixed with 1/3 volume of the Lenti-X concentrator. After incubation at 4 °C for 30 min, the mixture was centrifuged at 1,500 g at 4 °C for 45 min. The resulting pellet was resuspended in 50-100 µL of cell culture medium to infect target cells. Target cells were treated with 5 µg/mL polybrene (#TR-1003-G, Merck) shortly before infection. Successfully transduced cells were then positively selected by adding 2.5 µg/ml Puromycin to the medium. Expression of the constructs was induced by the addition of 1 μg/ml dox to the medium 24 hrs before probing the cells. Monoclonal cell lines were generated by limiting dilution or sorted for positive cells (BD FACS Aria Fusion II). Cells were screened for low background fluorescence via flow cytometry (BD LSRFortessa, BD FACSymphony). In brief, cells were detached, and single cells were strained through a cell strainer and suspended in flow buffer (PBS supplemented with 0.5% BSA, 2.5mM EDTA). Dead cells were excluded by DAPI staining and doublets were excluded using FCS/SSC gating.

### Cellular micropatterning on glass

To ensure consistency in the geometry and allow for cell shape control, most experiments were performed in micropatterned cells. 3T3 fibroblasts were seeded on 20 µm adhesive spots on glass dishes or 24-well glass-bottom plates. Adhesive spots were generated using an inverted Ti2^®^ Nikon microscope equipped with the Primo micropatterning module (Alvéole). In brief, 35 mm glass-bottom dishes (Greiner bio-one, 627860) were plasma treated for 1 minute (Gala Instrument, Plasma Prep 2) and a small PDMS stencil with an inner hole of ∼10-20 µL was placed in the middle of the dish to reduce reagent usage. Next, the glass surface inside the stencil was coated with a non-adhesive layer of 0.2 mg/mL polyethylene glycol (PLLg-PEG, Susos AG) for 1 hour at RT. Then, the surface was washed three times with PBS before applying a thin layer of 4-benzoylbenzyl-trimethylammonium chloride micropatterning reagent (PLPP, Alvéole). The sample was placed on the stage and the desired geometry (20 µm circles spaced 40 µm apart) was photo-patterned using a 375 nm laser (3.36 mW) to apply a laser dose of 2000 mJ/mm2, using the µmanager software v.1.4.22 with the Leonardo plugin v.4.12 (Alvéole). Finally, the surface was coated with 50 µg/mL bovine fibronectin (Merck, F1141) for 30 minutes and washed 3 times with PBS, before being stored at 4C for up to one week for experiments. For all experiments, 10^5^ cells were seeded onto the patterned area after detachment. Cells were allowed to settle and adhere for 20 minutes before non-attached cells were washed out. Experiments were performed 2 hours after cell seeding if not indicated otherwise.

### Dynamic micropatterning

For dynamic micropatterning, the steps for photo-patterning were the same as described above, but instead of coating with PLL-g-PEG, the surface is coated with 0.2 mg/ml of PLL(20)-g[3.5]- PEG(2)/PEG(3.4)-Biotin(50%) [SUSO, SZ44-79] after plasma treatment. After washing, 105 cells were seeded onto the patterned dish. Cells were allowed to settle and adhere for 20 minutes before non-attached cells were washed out. To release cells, conjugated streptavidin- fibronectin (see below) pre-diluted in media was added to cover the glass surface at a density of 311 ng/cm2.

Fibronectin was conjugated with streptavidin to enable binding to the PLL-g-PEG-Biotin. This was accomplished using a commercially available streptavidin conjugation kit (Streptavidin conjugation kit – Lightning Link, ab102921). Manufacturer’s instructions were followed for the conjugation of 100 µl (1 mg/ml) streptavidin-fibronectin. Briefly, 10 µl of modifier was added to 100 µl of fibronectin. The solution was mixed with a single lyophilized vial and left overnight at room temperature in the dark. The next day, 10 µl of quencher were added and after 30 minutes the streptavidin-fibronectin was ready for use,and stored at 4C.

### Drug treatments

NIH-3T3 fibroblasts were seeded on micropatterned dishes as described above and actomyosin- perturbing drugs were added directly to the media for a final concentration of 10 nM for latrunculin B (Merck, 428020), and 20-50 µM for para-amino-blebbistatin (Cayman Chemical, 2097734-03-5). Cells were incubated with the drugs for at least 30 minutes before image acquisition, dynamic micropatterning or magnetic pincher; the drugs were kept in the media during these experiments.

### Real time deformability cytometry

RT-DC experiments were performed in a previously described experimental setup^44^. Briefly, a poly(dimethylsiloxane) (PDMS; Sylgard 184, VWR) microfluidic chip is microfabricated with an inlet and outlet of 1.5 mm and two reservoirs connected by a 300-μm-long narrow channel (constriction) with a 30 μm × 30 μm square cross-section. The chip is then assembled on the xy stage an Axiovert 200M inverted microscope (Zeiss) and connected to a syringe pump (NemeSyS, Cetoni) for driving a cell suspension through the channel. Images of deformed cells were acquired using a CMOS camera (MC1362, Mikrotron). On the day of the experiment, NIH-3T3 cells were detached with 0.05% Trypsin-EDTA and suspended in medium supplemented with 0.5% methylcellulose and EDTA up to 15 mM. The latter was used to achieve two different osmolarities: 380 Osm and 290 Osm. NIH-3T3 were deformed using the 30 μm channel at two different flow rates: 0.16 and 0.24 μl/s. Analysis of deformation was performed as previous described, deriving the hydrodynamic stresses on the object surface and quantifying its deformation on the basis of linear-elasticity theory. This allows to decouple the relationship between cell size–dependent stress, deformation and elastic material parameters. Data was analysed with the ShapeOut software with an area ratio of 1-1.075, an area of 100-500 μm, an aspect ratio of 1-3, and a fluorescence range of 300-40000.

### Magnetic pincher experiments

#### Sample preparation, setup, and optogenetic activation

Superparamagnetic beads (Dynabeads M-450 Epoxy, ThermoFisher) were coated with a fresh solution of fibronectin 0.1 mg/mL in 100mM HEPES. Superparamagnetic beads coated with streptavidin (Dynabeads Streptavidin-Coated M-450, Thermo Fisher) were coated in a fresh solution of mPEG-(5K)- Biotin in 100 mM HEPES. Fibronectin-coated beads were stored at 4℃ for up to one week, while PEG-coated beads were stored up at 4℃ for up to four weeks. 24-48 hrs before magnetic pinching experiments, cells were seeded at a density of 5*10^3^ cells/cm2 in the presence of fibronectin-coated beads at a density of 10^4^ beads/cm2, for the cells to engulf them. On the day of the experiment, cells were detached from the substrate using 0.05% Trypsin-EDTA (Thermo Fisher Scientific) and plated on micropatterned dishes as described above. Then 1.5*10^5^ of PEG-coated beads were added to the dish, and the medium was supplemented by 20 mM of HEPES buffer, to mitigate pH fluctuations during the experiment.

All magnetic pincher experiments were performed on a basic, inverted microscope (Zeiss, Axio 164 Observer). Images were captured with a digital CMOS camera (Hamamatsu ORCA-Flash4.0, C11440). The microscope is equipped with the piezo-controller for the focus (PIFOC P-721.CDQ), and a 100X objective (Zeiss, Plan-Apchromatic, Aperture = 1.4). Activation on this microscope was performed using a fluorescent light source (Zeiss, HXP 120C), passing through a filter cube to visualize GFP (Zeiss, Filter 09, BP 450-490). The magnetic field is generated by two coaxial coils (SBEA) with mu metal core (length, 40 mm; diameter, 26 to 88 418 mm; 750 spires). The coils are powered by a bipolar operational power supply amplifier 6A/36V (Kepco) controlled by a data acquisition module (National Instruments).

The maximum field generated is 54 mT with a gradient less than 0.1 mT/mm over the sample. The setup was heated at 37° C, without CO2 supplementation and is controlled by a custom LabVIEW interface that ensures the synchronicity between piezo position, magnetic field imposition, and image acquisition. A resting field of 5 mT is maintained to hold a pincher across the cortex of a cell, without which the beads would freely float away. The experiment begins with a “passive” phase where triplet stacks spaced 500 nm in the z-axis are captured every 600 ms for a total duration of 8 seconds. This phase is used to derive an accurate estimation of the 3D distance between the beads to later calculate the thickness. Then, the magnetic field is tuned to execute compressive ramps. A compression is divided into 2 “action phases”: a descending sigmoid to a lower force 1(mT) which is maintained (constant phase) for two seconds to effectively “relax” the cortex. The field is then adjusted from 1 mT to 50 mT with a t4 functional form within 1.5 seconds, rapidly compressing the cortex. Images are captured with a 5 ms exposure time. The acquisition frequency varies between the phases, but is highest during the compression phase, set at 10-14 Hz. Only the mid-plane is imaged during the indentation to enable high- frequency acquisition. A total of 6 compression/constant field cycles were acquired for the baseline while 10 cycles were acquired for the activation phase.

Optogenetic activation was performed with a first long-exposure activation of 5 seconds, followed by pulses of 150 ms exposure time, at 20 s intervals (every compression). HXP lamp (HXP 120 C) was set to an intensity level 2 and the neutral density filter to 5. The resultant intensity is 0.3 mW/mm^2^. Filter set 09 (Zeiss) was used, giving an excitation wavelength of BP 450-490 nm. To avoid unwanted activation of optogenetic constructs with transmission light, a long-pass coloured filter (Thorlabs, FGL560S) was placed after the light source to block wavelengths below 560 nm.

#### Data analysis

Initially, the center of mass (XM-YM) of the beads was detected with a subpixel resolution using the “Analyze particles” function on Fiji. The resulting file is processed using a home-made tracking algorithm written in Python (https://github.com/jvermeil-biophys/CortExplore) which detects the trajectories in 3D of the centers of the two beads pinching the cortex. A depthograph, obtained from a 0.05 z stack for several beads, is used to determine the correct z position of each bead at each timelapse point. To obtain the thickness of the cortical layer, the 3D distance between the two beads is subtracted by the bead diameter. The pinching force was also computed knowing the distance between the beads, the external magnetic field and the magnetization function of the beads. To characterize the cortex thickness, the median of the cortical thickness at 1 mT (undeformed conditions) and 5 mT (nominal field) was measured. The elastic modulus at various stress ranges (50-200pN, 400-800pN, 500 pN) was computed from the strain-stress curve corresponding to each compression using a model developed by Chadwick for this specific geometry. Stiffness-thickness correlations were performed on individual cells for specific ranges to extract a normalized elastic modulus for a reference thickness.

### Simulations of contractile actin networks

Simulations proceeded in two phases: an equilibration phase followed by a compression of the simulation (**Fig. 2G**). During equilibration the system is allowed to reach a steady state over 22 million time-integration steps. The compression proceeds by a linear decrease of the z-direction from L_z=19 to L_z=9.5 over 2 million time-steps, corresponding to a linear increase in strain from 0 to 0.5.

To disentangle effects on the mechanics from the different components of the system, for every linker length we performed 4 different independent simulations: With linker and myosin binding turned on during equilibration and compression and with either linker or myosin binding disabled, as well as both actin and myosin binding disabled.

Simulations were performed by overdamped molecular dynamics in LAMMPS^100^ with the use of the REACTER package^101^ and results are visualized in OVITO^102^. Each actin filament was modelled as a polymer of 50 spherical beads, connected by the harmonic bond potential U_b(r)=K_b(r−r₀)² with equilibrium length r₀=a_s and harmonic constant K_b=500 e_s/a_s², where a_s is the simulation length unit and e_s=k_BT is the simulation energy unit, with k_B the Boltzmann constant and T the temperature, and by the harmonic angle potential U_θ(θ)=K_θ(θ−θ₀)² with K_θ=500 e_s rad⁻² and θ₀=π, leading to stiff filaments. Actin filaments were therefore approximately inextensible and fairly rigid, with a persistence length 20 times larger than the filament length; the knowledge of the persistence length of actin^103^ allows us to map the filament length to order ∼ 1 μm, and therefore the diameter of a single bead (a_s) to order ∼ 20 nm. a_s=1 is the simulation length unit, e_s=k_BT is the energy unit, and t_s=a_s√(m/e_s) is the time unit for bead mass m=1.

Each linker was modelled as a polymer of spherical beads of diameter a_s, connected by a harmonic bond of equilibrium length r₀=a_s and harmonic constant K_b=500 e_s/a_s². Linkers were therefore hard to stretch and floppy with elongated lengths 5a_S, 9a_s and 19a_s.

Each myosin unit was modelled as a dumbbell of 2 spherical beads of diameter a_s, connected by a spring of equilibrium length r₀=10a_s and harmonic constant K_b=20 e_s/a_s².

Volume exclusion between beads was modeled through a Weeks-Chandler-Anderson (WCA) potential of amplitude ε=10e_s and contact range a_s. For particle–particle exclusion, the LAMMPS cosine/squared WCA convention was used with σ_WCA=r_c=a_s and U_WCA(r)=10e_s[(a_s/r)¹²−2(a_s/r)⁶+1] for r<a_s, with U_WCA=0 for r≥a_s. The hard wall in z-direction instead uses the conventional Lennard-Jones parameter σ_LJ,wall=2⁻¹ᐟ⁶a_s=0.890899a_s with cutoff r_c=a_s; both conventions therefore give the same exclusion distance a_s.

#### Time integration

The dynamics resulted from a velocity-Verlet integrator of the equations of motion, coupled with a Langevin thermostat, such that the simulated ensemble was a priori NVT. The Langevin thermostat adds a viscous drag force and a random force to the force originating from inter-particle interactions, thus implicitly representing collisions with the solvent. The viscous drag force is F_drag=−(m/t_d)v, where v is the particle velocity and t_d is the characteristic damping time of the thermostat; 1/t_d can therefore be interpreted as an inverse reduced viscosity.

The characteristic damping time of the thermostat was set to t_d=0.1t_s. The integration timestep was Δt=0.01t_s and proved sufficient for numerical stability in the present systems.

#### Stress measurement

The forces exerted by the system on the top and bottom wall via the repulsive potential were collected and averaged over 25000 consecutive timesteps to obtain the stress-strain curves in (Fig 2 H) The differential modulus K was obtained by linear fits to constant width and overlapping stress intervals of the respective stress-strain relation (**Fig. 2I**). The simulations were performed with 3 different seeds in the random number generation for time-integration for the equilibration phase up to 20 million steps. These 3 steady states where then further integrated for 2 million steps at constant box height and 2 million steps of compression using 2 further seeds each. Hence, in total the stress-strain curves are average over 6 simulations.

#### Simulation box and densities

The simulation box was chosen to be anisotropic with box dimensions Lx=100a_s, Ly=100a_s and Lz=19a_s, having periodic boundary conditions in x- and y-direction. The upper and lower boundaries in z were hard walls with the short-ranged, purely repulsive Lennard-Jones potential described above. The system is comprised of 200 actin filaments, 300 myosins and 250 linkers. The actin filaments are initially places oriented along the xy-plane with random orientation and position in z. The myosins and linkers are placed randomly and uniformly in the box. Before the actual simulation a short time-integration was performed to resolve initial particle overlaps and to ensure numerical stability. Linker and myosin binding dynamics.

From the beginning of the simulation, we allowed linker tips to bind to unbound actin beads at the nominal rate k_on=1t_s⁻¹ for any two such beads in contact (implemented as probability p_on=0.1 every 10 timesteps). The formed bond was harmonic, with equilibrium length r₀=1.2a_s and strength K_b=80 e_s/a_s². Linker unbinding (bond deletion) occurred at the nominal rate k_off=2×10⁻⁵t_s⁻¹ (parameterized as p_off=2×10⁻⁶ every 10 timesteps and implemented equivalently as p=0.1 every 500,000 timesteps). Myosin beads could bind unbound actin beads with the same rate k_on=1t_s⁻¹ and with bond parameters r₀=1.2a_s and K_b=20 e_s/a_s². A myosin bead would walk along an actin filament by moving the bond that connects it to an actin bead to the next actin bead, in the direction of the barbed end. This occurred at rate k_walk=10t_s⁻¹ (probability 1 every 10 timesteps), provided that the next actin bead was not already bound to another myosin and was within distance 1.5a_s.

### LED array for cell illumination

The LED array used for activation and immunostaining experiments was manufactured by Alejandro Gil Ortiz at the EMBL Electronics Workshop. It consists of a PCB control board connected to an LED board of PLCC4 Surface Mount LEDs (OVSA1xBC2R8 Series) with a wavelength of 470nm and arranged in the same layout as a 24-well plate. The control board can be plugged to the serial port of any laptop and configured through a custom Python script using the pyserial library.

### Optogenetic activation and immunofluorescence

Cells were seeded on micropatterned glass-bottom, dark-coloured 24-well plates (reference) as described in the “micropatterning” section and activated with the LED array for 2 minutes with three sets of pulses (1.5s ON, 0.5s OFF) intercalated by 20s of dark (recovery time). 2 LEDs at 75% of their maximum intensity were used for illumination. After activation with LED array, cells were fixed and permeabilized simultaneously by adding prewarmed (37 ° C) fixation buffer (2x) to growth medium and incubated for 20 min at RT. Fixation buffer (1x) contains 4% PFA (Thermo Fisher Scientific), 0.5% Tergitol (Serva), 20mM sucrose (Merck) in cytoskeleton buffer (10 mM MES, 140 mM KCl, 3 mM MgCl2, 2 mM EGTA, pH = 7.5). Cells were washed trice for 10 min with TBS (Tris-Buffered Saline) and stained with CellBrite Blue (Biotium, #30024) to label the membrane. Cells were incubated for 12 minutes at RT in the dark with CellBrite Loading Dye and Imaging Solution (1:100 in TBS) and washed trice with TBS. Subsequently, cells were blocked for 1 hour with 5% BSA (Merck) in TBS. Primary antibody staining was performed over night at 4 ° C in 1% BSA in TBS (anti-Phospho-MLC2 (Ser19), #3671, Cell signaling technology, 1:150). Cells were washed four times over 20 minutes with TBS and Secondary antibody staining was performed for 1 hour at RT in 1% BSA in TBS (AF647 anti-rabbit #A21244, Thermo Fisher Scientific, 1:1.000). After washing thrice with TBS, cells were incubated with 500 uL of AF488 Phalloidin (#A12379, Thermo Fisher Scientific, 1:1.000) at 4 ° C until imaging.

Cells were imaged using an inverted confocal field scanning spinning disk and high-resolution Nikon-CSU-W1 SORA microscope equipped with a scMOS camera (Hamamatsu Orca FusionBT), four lasers (405 nm, 488 nm, 561 nm, 647 nm), and a 60X Water objective with an NA of 1.27, controlled by NIS-Elements software. Cells were imaged by acquiring z-stacks of 0.5 µm with a SoRA 4x magnifier.

### Fluorescence microscopy and optogenetic activation in live cells

Live fluorescence imaging and optogenetic activation was performed on a Zeiss LSM980 Airyscan microscope equipped with four diode lasers (405nm, 488nm, 561nm, 639nm), fast live-SR acquisition mode, and a 63x Oil objective with an NA of 1.4, controlled by ZenBlue software. Cells were imaged live at 37°C and 5% CO_2_ after being seeded onto micropatterns 2-4 hours before imaging. A feedback microscopy routine was written with ZenBlue macro language and AutoMicTools to enable automated autofocus and multiposition live imaging coupled to optogenetic activation. For each position, autofocus was determined using the reflective xz image and a Fiji script, after which a fixed focal plane of 5 µm was set for subsequent acquisitions. Cells were then imaged by acquiring a z-stack of 0.5 µm followed by a baseline phase of dual-channel acquisitions with images taken every 10s. Cells were subsequently segmented on-the-fly with Fiji, and the segmentation mask was overlaid onto ZenBlue to determine the activation region. Cells were activated every 60s with a 300ms pulse of 350W/m2 (0.1% intensity) using the “Bleaching” function of the ZenBlue software, while coupled to a timelapse taken every 10s. A second z-stack of 0.5 µm was taken after activation. Prior to image acquisition, the cells were incubated with 1:1000 SPY650-FastAct (SC505, Spirochrome) diluted in DMSO (as per manufacturer’s instructions) for 1 hour.

### Dynamic micropatterning and optogenetic activation

Dynamic micropatterning experiments in live cells were performed using a Nikon Ti-E inverted widefield microscope equipped with a SpectraX light source. (Lumencor), a Hammamatsu Orca Flash 4 V2, and a 20x Air phase contrast objective (Nikon PH2) with a NA of 0.75. Cells were imaged live at 37°C and 5% CO_2_ after being seeded onto micropatterns 2-4 hours before imaging. Phase contrast images were taken every 2min at 5 different fields of view (FOVs) for three different phases: baseline, activation, and post-release. Activation pulses lasted 700ms and were delivered using the GFP excitation filter (485/20) at an intensity of 5%; for slow activation protocols, 1 pulse was delivered every 1 or 2 min, while for fast activation protocols 1 pulse was delivered every 20 s. After activation, 10 µL of Fibronectin-SA (pre-diluted in 100µL of media) were added to the dish to release the cells from the micropatterns. Cells were recorded for a maximum of 50 minutes after release from micropatterns. To avoid unwanted activation of optogenetic constructs with transmission light throughout the entire experiment, a long-pass coloured filter (Edmund Optics) was placed after the light source to block wavelengths below 560nm.

### Image Analysis

#### Actin/phospho-myosin mean fluorescence intensity

z-stacks of cells were acquired in 4 channels: our expressed constructs, phalloidin, myosin, and a membrane stain. Cell masks were generated in Ilastik for the CellBrite 405 membrane stain (pixel classification, probability maps) and fluorescence intensities were measured using a custom-made Fiji macro. Briefly, cell masks were eroded and dilated to generate a band encompassing the membrane stain and therefore the cortical region. Mean actin and phospho-myosin fluorescence intensity was measured across the z-slices where this cortical band could be defined. For each condition, mean fluorescence intensities were normalized to the average of the control condition for each replicate.

#### Polarization and delamination analysis

Cell masks were generated in Ilastik for the actin channel (pixel classification, probability maps) and loaded per frame together with the raw fluorescence image stack for subsequent processing in Python using the skimage and scipy packages.

#### Delamination analysis

The resulting masks were eroded by 3 pixels to avoid membrane-boundary segmentation artefacts. For each frame, a Euclidean distance transform was computed inside the eroded mask, giving each interior pixel its distance from the nearest point on the cell boundary. The interior was then divided into 15 equal- width concentric shells (band 0 = outermost/cortical) spanning 0 to the maximum in-mask distance, and the mean fluorescence intensity was computed within each shell. The mean actin intensity of the 2 outermost bands (cortical) was divided by the mean intensity of the 3 following bands (subcortical):

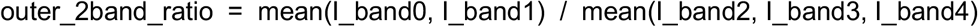

This outer/inner-band ratio was used as a measurement of cortical actin enrichment, and decreases as the cortex delaminates from the membrane.

#### Polarity index analysis

The resulting masks were eroded by 15 pixels to restrict the analysis to a peripheral cortical band, and the inner boundary of the eroded mask was extracted and dilated by 3px to give a finite-width sampling band. The cell outline was traced, and the raw fluorescence intensity was sampled at each contour point. The resulting 1-D intensity trace was smoothed (moving average, window = 3 points) and linearly interpolated onto a fixed grid of 360 points, giving one intensity value per degree of angular position around the cell periphery. Kymographs were obtained by stacking this angular intensity profile over all frames.

For each frame, the angular intensity profile I(θ) (θ = 0–359°) was treated as a circular weighted distribution. The weighted circular mean defined the instantaneous polarization center:

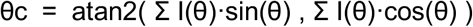

The degree of angular clustering was quantified by the mean resultant length R:

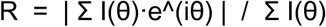

R ranges from 0 (intensity uniformly distributed around the periphery, no polarization) to 1 (intensity fully concentrated at a single angle, maximal polarization).

A local enrichment metric, ROI30, was computed as the fraction of total peripheral intensity falling within a ±30° window (60° total) centered on θc:

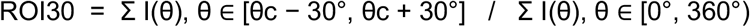

The composite polarity index (PI) combines both terms:

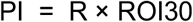

Therefore, a high PI requires both a concentrated (high-R) and locally confined (high-ROI30) distribution of signal, distinguishing true focal polarization from broad or diffuse asymmetries. Baseline R (mean of the first 24 frames, pre-activation) was used to compute fold-change and threshold- based (ΔR ≥ 0.1) onset detection of polarization.

#### Spreading directionality analysis

For 3D segmentation of phase contrast timelapses of spreading cells, we trained a vision foundation model using Micro-SAM ^104^ and torch-em ^105^ with manually curated phase contrast segmentations obtained in Micro-SAM. Cell outlines were provided as binary segmentation-mask time-lapses and processed on Python using the numpy package. A cell was excluded from analysis if more than 3 of its first 10 frames contained an empty mask, indicating segmentation or tracking failure at the start of the time-lapse.

A reference center (cy, cx) and initial radius R0 were defined by fitting a circle to the boundary of the first non-empty mask (Kasa algebraic circle fit); the boundary was defined as foreground pixels with at least one background neighbor. As alternatives, R0 could instead be set to the median distance from the centroid to the boundary, or to the radius of a circle with the same area as the mask. For each frame, the mask was sampled in 72 angular bins (5° each) around the reference center (cy, cx). In each bin, the maximum distance from the center to the cell edge was recorded, giving R(t, θ) — how far the edge sits from the center, in that direction, at time t. Radial expansion relative to the starting geometry was then given by:

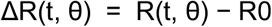

Only outward growth was counted as spreading — frame-to-frame retractions were ignored — and these outward steps were accumulated over time for each angle:

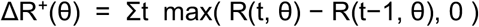

This cumulative outward-growth profile was treated like a weighted compass: each angle θ contributes a vector of length ΔR⁺(θ), and their weighted average (a circular mean) gives both an overall spreading direction, θ_spread, and a directionality score, |V_spread| — 0 for spreading equally in all directions, 1 for spreading confined to a single direction. Repeating this calculation cumulatively at every frame, instead of only at the end, yielded a Directionality Index time-course, DI(t): how directional the spreading has been up to that point, independent of how much the cell has spread overall.

Spreading onset was defined as the first frame at which the mean radial expansion (mean ΔR(t, θ) across all angles) exceeded 5 px and remained above this threshold in the following frames, rather than reverting immediately. Onset was then used as the reference point for evaluating directionality over two time-windows: an early-spreading window (the 10 minutes following onset) and the DI(t) time-course over the full acquisition (50 minutes).

## Code availability

The Fiji macro for quantifying cortical actin and the Python scripts for quantifying cortical dynamics and directional cell spreading will made available upon publication at https://github.com/Fnunesv.

## Statistical analysis

Statistical analyses are described in all figure legends and were performed using R. Data visualization was done using R. Normality of data distribution was tested by Shapiro-Wilk test. For magnetic pincher data, a paired two-tailed t-test was used for normally distributed data when only 2 conditions had to be compared. Otherwise, a non-parametric Wilcox signed-rank test was used.

To account for the nested structure of cortical fluorescence intensity and cell spreading dynamics (multiple cells measured within each independent experiment), differences between conditions were assessed using beta generalized linear mixed models (beta-GLMM) ^106^ . Pairwise condition contrasts were obtained from estimated marginal means using the emmeans R package, and p-values across the set of planned comparisons were adjusted for multiple testing with the Holm-Bonferroni method.

In all box plots, the lower and upper hinges correspond to the first and third quartiles (the 25th and 75th percentiles). Black/solid line corresponds to the median. In violin plots the colored area reflects the probability distribution of the data points.

## REFERENCES

1. Levayer, R. & Lecuit, T. Biomechanical regulation of contractility: spatial control and dynamics. Trends in Cell Biology 22, 61–81 (2012).

2. Salbreux, G., Charras, G. & Paluch, E. Actin cortex mechanics and cellular morphogenesis. Trends in Cell Biology 22, 536–545 (2012).

3. Chugh, P. & Paluch, E. K. The actin cortex at a glance. Journal of Cell Science 131, jcs186254 (2018).

4. Banerjee, S., Gardel, M. L. & Schwarz, U. S. The Actin Cytoskeleton as an Active Adaptive Material. Annu. Rev. Condens. Matter Phys. 11, 421–439 (2020).

5. Murrell, M., Oakes, P. W., Lenz, M. & Gardel, M. L. Forcing cells into shape: the mechanics of actomyosin contractility. Nature Reviews Molecular Cell Biology 16, 486–498 (2015).

6. Koenderink, G. H. & Paluch, E. K. Architecture shapes contractility in actomyosin networks. Current Opinion in Cell Biology 50, 79–85 (2018).

7. Sobral, A. F. et al. Plastin and spectrin cooperate to stabilize the actomyosin cortex during cytokinesis. Current Biology 31, 5415–5428.e10 (2021).

8. Toyoda, Y. et al. Genome-scale single-cell mechanical phenotyping reveals disease-related genes involved in mitotic rounding. Nat Commun 8, 1266 (2017).

9. Ding, W. Y. et al. Plastin increases cortical connectivity to facilitate robust polarization and timely cytokinesis. Journal of Cell Biology 216, 1371–1386 (2017).

10. Taneja, N. et al. The actin crosslinker Fimbrin is required for force sensing at tricellular junctions. Developmental Cell 61, 1699–1716.e12 (2026).

11. Paradžik, T., Podgorski, I. I., Vojvoda Zeljko, T. & Paradžik, M. Ancient Origins of Cytoskeletal Crosstalk: Spectraplakin-like Proteins Precede the Emergence of Cortical Microtubule Stabilization Complexes as Crosslinkers. IJMS 23, 5594 (2022).

12. Sedeh, R. S. et al. Structure, Evolutionary Conservation, and Conformational Dynamics of Homo sapiens Fascin-1, an F-actin Crosslinking Protein. Journal of Molecular Biology 400, 589–604 (2010).

13. Liem, R. K. H. Cytoskeletal Integrators: The Spectrin Superfamily. Cold Spring Harb Perspect Biol 8, a018259 (2016).

14. Gong, R. et al. Fascin structural plasticity mediates flexible actin bundle construction. Nat Struct Mol Biol 32, 940–952 (2025).

15. Alvarado, J., Sheinman, M., Sharma, A., MacKintosh, F. C. & Koenderink, G. H. Molecular motors robustly drive active gels to a critically connected state. Nature Phys 9, 591–597 (2013).

16. Ennomani, H. et al. Architecture and Connectivity Govern Actin Network Contractility. Current Biology 26, 616–626 (2016).

17. Chaudhuri, O., Parekh, S. H. & Fletcher, D. A. Reversible stress softening of actin networks. Nature 445, 295–298 (2007).

18. Gardel, M. L. et al. Elastic Behavior of Cross-Linked and Bundled Actin Networks. Science 304, 1301– 1305 (2004).

19. Jansen, S. et al. Mechanism of Actin Filament Bundling by Fascin. Journal of Biological Chemistry 286, 30087–30096 (2011).

20. Mei, L. et al. Structural mechanism for bidirectional actin cross-linking by T-plastin. Proceedings of the National Academy of Sciences 119, e2205370119 (2022).

21. Meyer, R. K. & Aebi, U. Bundling of actin filaments by alpha-actinin depends on its molecular length. J Cell Biol 110, 2013–2024 (1990).

22. Wachsstock, D. H., Schwartz, W. H. & Pollard, T. D. Affinity of alpha-actinin for actin determines the structure and mechanical properties of actin filament gels. Biophysical Journal 65, 205–214 (1993).

23. Winkelman, J. D. et al. Fascin- and α-Actinin-Bundled Networks Contain Intrinsic Structural Features that Drive Protein Sorting. Current Biology 26, 2697–2706 (2016).

24. Gorlin, J. B. et al. Human endothelial actin-binding protein (ABP-280, nonmuscle filamin): a molecular leaf spring. Journal of Cell Biology 111, 1089–1105 (1990).

25. Schwaiger, I., Kardinal, A., Schleicher, M., Noegel, A. A. & Rief, M. A mechanical unfolding intermediate in an actin-crosslinking protein. Nature Structural & Molecular Biology 11, 81–85 (2004).

26. Pan, L., Yan, R., Li, W. & Xu, K. Super-Resolution Microscopy Reveals the Native Ultrastructure of the Erythrocyte Cytoskeleton. Cell Reports 22, 1151–1158 (2018).

27. Xu, K., Zhong, G. & Zhuang, X. Actin, Spectrin, and Associated Proteins Form a Periodic Cytoskeletal Structure in Axons. Science 339, 452–456 (2013).

28. Kumar, A. et al. Filamin A mediates isotropic distribution of applied force across the actin network. Journal of Cell Biology 218, 2481–2491 (2019).

29. Scheff, D. R. et al. Actin filament alignment causes mechanical hysteresis in cross-linked networks. Soft Matter 17, 5499–5507 (2021).

30. Freedman, S. L. et al. Mechanical and kinetic factors drive sorting of F-actin cross-linkers on bundles. Proc. Natl. Acad. Sci. U.S.A. 116, 16192–16197 (2019).

31. Sakamoto, R. & Murrell, M. P. F-actin architecture determines the conversion of chemical energy into mechanical work. Nature Communications 15, 3444 (2024).

32. Sakamoto, R., Sun, Z. G. & Murrell, M. P. Crosslinked F-actin networks regulate load-dependent energy conversion. Commun Biol 9, 572 (2026).

33. Weirich, K. L., Stam, S., Munro, E. & Gardel, M. L. Actin bundle architecture and mechanics regulate myosin II force generation. Biophysical Journal 120, 1957–1970 (2021).

34. Lazarides, E. & Burridge, K. α-Actinin: Immunofluorescent localization of a muscle structural protein in nonmuscle cells. Cell 6, 289–298 (1975).

35. Bretscher, A. & Weber, K. Fimbrin, a new microfilament-associated protein present in microvilli and other cell surface structures. The Journal of cell biology 86, 335–340 (1980).

36. Krueger, D., Quinkler, T., Mortensen, S. A., Sachse, C. & De Renzis, S. Cross-linker–mediated regulation of actin network organization controls tissue morphogenesis. Journal of Cell Biology 218, 2743– 2761 (2019).

37. He, J. et al. Prevalent presence of periodic actin–spectrin-based membrane skeleton in a broad range of neuronal cell types and animal species. Proceedings of the National Academy of Sciences 113, 6029–6034 (2016).

38. Huang, C. Y. M. et al. αII spectrin forms a periodic cytoskeleton at the axon initial segment and is required for nervous system function. Journal of Neuroscience 37, 11311–11322 (2017).

39. Mahlandt, E. K. et al. Opto-RhoGEFs: an optimized optogenetic toolbox to reversibly control Rho GTPase activity on a global to subcellular scale, enabling precise control over vascular endothelial barrier strength. Preprint at 10.7554/eLife.84364.1 (2023).

40. Martínez-Ara, G. et al. Optogenetic control of apical constriction induces synthetic morphogenesis in mammalian tissues. Nat Commun 13, 5400 (2022).

41. Yamamoto, K. et al. Optogenetic relaxation of actomyosin contractility uncovers mechanistic roles of cortical tension during cytokinesis. Nat Commun 12, 7145 (2021).

42. Park, H. et al. Optogenetic protein clustering through fluorescent protein tagging and extension of CRY2. Nat Commun 8, 30 (2017).

43. Taslimi, A. et al. An optimized optogenetic clustering tool for probing protein interaction and function. Nat Commun 5, 4925 (2014).

44. Otto, O. et al. Real-time deformability cytometry: on-the-fly cell mechanical phenotyping. Nat Methods 12, 199–202 (2015).

45. Jacobs, S. A. et al. Design of novel FN3 domains with high stability by a consensus sequence approach. Protein Engineering Design and Selection 25, 107–117 (2012).

46. Son, S. et al. Molecular height measurement by cell surface optical profilometry (CSOP). Proc. Natl. Acad. Sci. U.S.A. 117, 14209–14219 (2020).

47. Takatori, S. C., Son, S., Lee, D. S. W. & Fletcher, D. A. Engineered molecular sensors for quantifying cell surface crowding. Proc. Natl. Acad. Sci. U.S.A. 120, e2219778120 (2023).

48. Bakalar, M. H. et al. Size-Dependent Segregation Controls Macrophage Phagocytosis of Antibody- Opsonized Targets. Cell 174, 131–142.e13 (2018).

49. Abramson, J. et al. Accurate structure prediction of biomolecular interactions with AlphaFold 3. Nature 630, 493–500 (2024).

50. Mokbel, M. et al. Numerical Simulation of Real-Time Deformability Cytometry To Extract Cell Mechanical Properties. ACS Biomater. Sci. Eng. 3, 2962–2973 (2017).

51. Descovich, C. P. et al. Cross-linkers both drive and brake cytoskeletal remodeling and furrowing in cytokinesis. MBoC 29, 622–631 (2018).

52. Chugh, P. et al. Actin cortex architecture regulates cell surface tension. Nature Cell Biology 19, 689– 697 (2017).

53. Laplaud, V. et al. Pinching the cortex of live cells reveals thickness instabilities caused by myosin II motors. SCIENCE ADVANCES (2021).

54. Lembo, S. et al. The distance between the plasma membrane and the actomyosin cortex acts as a nanogate to control cell surface mechanics. Preprint at 10.1101/2023.01.31.526409 (2023).

55. Vermeil, J. et al. A Magnetic Pincher for the Dynamic Measurement of the Actin Cortex Thickness in Live Cells. in Imaging Cell Signaling (eds Wuelfing, C. & Murphy, R. F.) 115–145 (Springer US, New York, NY, 2024). doi:10.1007/978-1-0716-3834-7_10.

56. Dar, S. et al. Caging of membrane-to-cortex attachment proteins can trigger cellular symmetry breaking. bioRxiv 10.1101/2024.10.14.618153 (2024) doi:10.1101/2024.10.14.618153.

57. Naganathan, S. R. et al. Morphogenetic degeneracies in the actomyosin cortex. eLife 7, e37677 (2018).

58. Fritzsche, M., Lewalle, A., Duke, T., Kruse, K. & Charras, G. Analysis of turnover dynamics of the submembranous actin cortex. MBoC 24, 757–767 (2013).

59. Wittmann, T., Dema, A. & Van Haren, J. Lights, cytoskeleton, action: Optogenetic control of cell dynamics. Current Opinion in Cell Biology 66, 1–10 (2020).

60. Yu, Q., Li, J., Murrell, M. P. & Kim, T. Balance between Force Generation and Relaxation Leads to Pulsed Contraction of Actomyosin Networks. Biophysical Journal 115, 2003–2013 (2018).

61. Kadzik, R. S., Maxian, O., Thomas, I., Kovar, D. R. & Munro, E. M. Rapid actin filament turnover maintains cortical connectivity while allowing for cell cortex deformation and flow.

62. Machacek, M. et al. Coordination of Rho GTPase activities during cell protrusion. Nature 461, 99–103 (2009).

63. Diz-Muñoz, A. et al. Membrane Tension Acts Through PLD2 and mTORC2 to Limit Actin Network Assembly During Neutrophil Migration. PLoS Biology 14, 1–30 (2016).

64. De Belly, H. & Weiner, O. D. Follow the flow: Actin and membrane act as an integrated system to globally coordinate cell shape and movement. Current Opinion in Cell Biology 89, 102392 (2024).

65. Isomursu, A. et al. Dynamic Micropatterning Reveals Substrate-Dependent Differences in the Geometric Control of Cell Polarization and Migration. Small Methods 8, 2300719 (2024).

66. Ebashi, S. & Ebashi, F. Alpha-actinin, a new structural protein from striated muscle. I. Preparation and action on actomyosinàtp interaction. J Biochem 58, 7–12 (1965).

67. Maruyama, K. & Ebashi, S. Alpha-actinin, a new structural protein from striated muscle. II. Action on actin. J Biochem 58, 13–19 (1965).

68. Gardel, M. L. et al. Elastic behavior of cross-linked and bundled actin networks. Science 304, 1301– 1305 (2004).

69. Reymann, A.-C. et al. Actin Network Architecture Can Determine Myosin Motor Activity. Science 336, 1310–1314 (2012).

70. Wagner, B., Tharmann, R., Haase, I., Fischer, M. & Bausch, A. R. Cytoskeletal polymer networks: The molecular structure of cross-linkers determines macroscopic properties. Proc. Natl. Acad. Sci. U.S.A. 103, 13974–13978 (2006).

71. Fritzsche, M., Erlenkämper, C., Moeendarbary, E., Charras, G. & Kruse, K. Actin kinetics shapes cortical network structure and mechanics. Sci. Adv. 2, e1501337 (2016).

72. Hosseini, K., Sbosny, L., Poser, I. & Fischer-Friedrich, E. Binding Dynamics of α-Actinin-4 in Dependence of Actin Cortex Tension. Biophysical Journal 119, 1091–1107 (2020).

73. Plaza, G. R., Uyeda, T. Q. P., Mirzaei, Z. & Simmons, C. A. Study of the influence of actin-binding proteins using linear analyses of cell deformability. Soft Matter 11, 5435–5446 (2015).

74. Laplaud, V., et al. Pinching the Cortex of Live Cells Reveals Thickness Instabilities Caused by Myosin II Motors. Sci. Adv vol. 7 http://advances.sciencemag.org/ (2021).

75. Truong Quang, B. A., et al. Extent of myosin penetration within the actin cortex regulates cell surface mechanics. Nat Commun 12, 6511 (2021).

76. Koenderink, G. H. et al. An active biopolymer network controlled by molecular motors. Proc. Natl. Acad. Sci. U.S.A. 106, 15192–15197 (2009).

77. Flormann, D. A. D. et al. The structure and mechanics of the cell cortex depend on the location and adhesion state. Proc. Natl. Acad. Sci. U.S.A. 121, e2320372121 (2024).

78. Broedersz, C. P., Storm, C. & MacKintosh, F. C. Nonlinear Elasticity of Composite Networks of Stiff Biopolymers with Flexible Linkers. Phys. Rev. Lett. 101, 118103 (2008).

79. Pogoda, K. et al. Unique Role of Vimentin Networks in Compression Stiffening of Cells and Protection of Nuclei from Compressive Stress. Nano Lett. 22, 4725–4732 (2022).

80. Golde, T. et al. Dynamics of supracellular keratin bundling and nuclear uncaging in stretched epithelia. Nature Physics 10.1038/s41567-026-03371-8 (2026) doi:10.1038/s41567-026-03371-8.

81. Bray, D. & White, J. G. Cortical Flow in Animal Cells. Science 239, 883–888 (1988).

82. Mayer, M., Depken, M., Bois, J. S., Jülicher, F. & Grill, S. W. Anisotropies in cortical tension reveal the physical basis of polarizing cortical flows. Nature 467, 617–621 (2010).

83. McFadden, W. M., McCall, P. M., Gardel, M. L. & Munro, E. M. Filament turnover tunes both force generation and dissipation to control long-range flows in a model actomyosin cortex. PLoS Comput Biol 13, e1005811 (2017).

84. Ding, W. Y. et al. Plastin increases cortical connectivity to facilitate robust polarization and timely cytokinesis. Journal of Cell Biology 216, 1371–1386 (2017).

85. Yolland, L. et al. Persistent and polarized global actin flow is essential for directionality during cell migration. Nat Cell Biol 21, 1370–1381 (2019).

86. Abu Shah, E. & Keren, K. Symmetry breaking in reconstituted actin cortices. eLife 3, e01433 (2014).

87. López-Gay, J. M. et al. Apical stress fibers enable a scaling between cell mechanical response and area in epithelial tissue. Science 370, eabb2169 (2020).

88. Van Der Gucht, J., Paluch, E., Plastino, J. & Sykes, C. Stress release drives symmetry breaking for actin-based movement. Proc. Natl. Acad. Sci. U.S.A. 102, 7847–7852 (2005).

89. Bement, W. M. et al. Activator–inhibitor coupling between Rho signalling and actin assembly makes the cell cortex an excitable medium. Nat Cell Biol 17, 1471–1483 (2015).

90. Munro, E., Nance, J. & Priess, J. R. Cortical Flows Powered by Asymmetrical Contraction Transport PAR Proteins to Establish and Maintain Anterior-Posterior Polarity in the Early C. elegans Embryo. Developmental Cell 7, 413–424 (2004).

91. Mak, M., Zaman, M. H., Kamm, R. D. & Kim, T. Interplay of active processes modulates tension and drives phase transition in self-renewing, motor-driven cytoskeletal networks. Nat Commun 7, 10323 (2016).

92. Driscoll, M. K., Losert, W., Jacobson, K. & Kapustina, M. Spatiotemporal relationships between the cell shape and the actomyosin cortex of periodically protruding cells. Cytoskeleton 72, 268–281 (2015).

93. Roth, H. et al. Filamin A promotes efficient migration and phagocytosis of neutrophil-like HL-60 cells. European Journal of Cell Biology 96, 553–566 (2017).

94. Ruprecht, V. et al. Cortical Contractility Triggers a Stochastic Switch to Fast Amoeboid Cell Motility. Cell 160, 673–685 (2015).

95. Lomakin, A. J. et al. Competition for actin between two distinct F-actin networks defines a bistable switch for cell polarization. Nat Cell Biol 17, 1435–1445 (2015).

96. Maiuri, P. et al. Actin Flows Mediate a Universal Coupling between Cell Speed and Cell Persistence Cell 161, 374–386 (2015).

97. De Belly, H. et al. Long-range mutual activation establishes Rho and Rac polarity during cell migration. Nat Cell Biol 28, 1244–1257 (2026).

98. Tichy, A.-M., Gerrard, E. J., Legrand, J. M. D., Hobbs, R. M. & Janovjak, H. Engineering Strategy and Vector Library for the Rapid Generation of Modular Light-Controlled Protein–Protein Interactions. Journal of Molecular Biology 431, 3046–3055 (2019).

99. Sladitschek, H. L. & Neveu, P. A. MXS-Chaining: A Highly Efficient Cloning Platform for Imaging and Flow Cytometry Approaches in Mammalian Systems. PLOS ONE 10, e0124958 (2015).

100. Thompson, A. P. et al. LAMMPS - a flexible simulation tool for particle-based materials modeling at the atomic, meso, and continuum scales. Computer Physics Communications 271, 108171 (2022).

101. Gissinger, J. R., Jensen, B. D. & Wise, K. E. REACTER: A Heuristic Method for Reactive Molecular Dynamics. Macromolecules 53, 9953–9961 (2020).

102. Stukowski, A. Visualization and analysis of atomistic simulation data with OVITO–the Open Visualization Tool. *Modelling Simul*. Mater. Sci. Eng. 18, 015012 (2010).

103. Ott, A., Magnasco, M., Simon, A. & Libchaber, A. Measurement of the persistence length of polymerized actin using fluorescence microscopy. *Phys*. Rev. E 48, R1642–R1645 (1993).

104. Archit, A. et al. Segment Anything for Microscopy. Nat Methods 22, 579–591 (2025).

105. Kirillov, A., et al. Segment Anything. Preprint at 10.48550/arXiv.2304.02643 (2023).

106. Brooks, M., E., et al. glmmTMB Balances Speed and Flexibility Among Packages for Zero-inflated Generalized Linear Mixed Modeling. The R Journal 9, 378 (2017).

